# How small molecules stabilize oligomers of a phase-separating disordered protein

**DOI:** 10.1101/2025.11.20.689623

**Authors:** Jiaqi Zhu, Thomas R. Sisk, Borja Mateos, Stasė Bielskutė-Garcia, Xavier Salvatella, Paul Robustelli

## Abstract

Small molecule inhibitors of the intrinsically disordered androgen receptor activation domain have been tested in clinical trials for the treatment of castration-resistant prostate cancer. These compounds have been shown to stabilize oligomeric forms of the androgen receptor activation domain in solution and alter the properties of androgen receptor condensates. The molecular mechanisms by which small molecules modulate these processes have been poorly understood in atomic detail. Here, we use long-timescale all-atom molecular dynamics (MD) simulations and nuclear magnetic resonance (NMR) spectroscopy to determine how small molecules stabilize highly dynamic, heterogeneous intermolecular interfaces that mediate oligomerization of the androgen receptor activation domain. The mechanisms determined here explain the relative potencies of androgen receptor activation domain inhibitors and suggest general strategies for designing small molecules that target oligomeric and, potentially, condensed forms of intrinsically disordered proteins.

---

Intrinsically disordered proteins (IDPs) play central roles in cellular signaling and regulation and are increasingly being pursued as drug targets^1–3^. The structural plasticity of IDPs enables them to form transient multivalent interactions that drive the assembly of biomolecular condensates^1,4^. This plasticity also makes IDPs extremely challenging drug targets. Several small molecules have been discovered that bind and inhibit specific IDPs and IDP-targeting compounds have entered human trials^3^. Biophysical experiments^5–12^ and atomistic molecular dynamics (MD) computer simulations ^11–20^ have shown that IDPs can remain disordered upon binding small molecules, and that the selectivity and affinity of IDP ligands discovered thus far is conferred through dynamic networks of transient interactions that may only subtly shift the conformational ensembles of IDPs.

The androgen receptor (AR) is a transcription factor containing an intrinsically disordered activation domain (AD)^21^. Small molecules targeting the disordered AR AD have shown promise for the treatment of castration-resistant prostate cancer (CRPC), as they inhibit AR splice variants lacking a ligand-binding domain that confer resistance to FDA-approved anti-androgen drugs^22,23^. The AR AD mediates the formation of transcriptional AR condensates in cells^12,24–26^. Nuclear magnetic resonance (NMR) spectroscopy, biophysical characterization of AR condensates, and cellular assays have identified aromatic residues and short transient helices in transactivation unit 5 (Tau-5) as the primary drivers of AR oligomerization and phase separation^12,26,27^. Tau-5 contains the interaction site of EPI-001^5,14,26^, a small molecule whose stereoisomer EPI-002 was developed into the prodrug EPI-506 for clinical testing in CRPC^3,22,23^. Recent studies showed that EPI-001 binds oligomeric Tau-5 with higher affinity than monomeric Tau-5, reduces the condensation cloud point (*T*_C_) of Tau-5, rigidifies AR condensates, and reduces the ability of AR condensates to recruit RNA polymerase II in live cells^12,26^. These observations raise a central mechanistic question: what molecular features of dynamic Tau-5 oligomerization interfaces are recognized and stabilized by AR AD inhibitors?

We previously employed all-atom MD simulations to determine atomic-resolution binding mechanisms of EPI-002, the most potent stereoisomer of EPI-001^5^, and related inhibitors to monomeric Tau-5^12,14,17^. These simulations, which are consistent with NMR measurements, showed that AR AD inhibitors predominantly interact with the partially helical R2 and R3 regions of Tau-5 and stabilize collapsed yet highly dynamic, molten-globule-like helical states. The AR AD inhibitor 1aa, which replaces the gem-dimethyl linker of EPI-002 with a linear alkyne (**Figure 1A**), has higher affinity for Tau-5 and more potently inhibits AR transcriptional activity in androgen-induced PSA-luciferase reporter assays^12^. A related diphenylacetylene derivative, 1ae, more effectively partitions into AR condensates, produces a larger reduction in *T*_C_, and has improved potency in cellular and human xenograft mouse models of CRPC relative to EPI-001^12^. Our previous monomer simulations suggest that the improved potency of 1aa and related compounds containing alkyne linkers may be associated with their ability to simultaneously intercalate into partially helical conformations of R2 and R3 while forming aromatic stacking interactions with residues in both regions.

**Figure 1:**
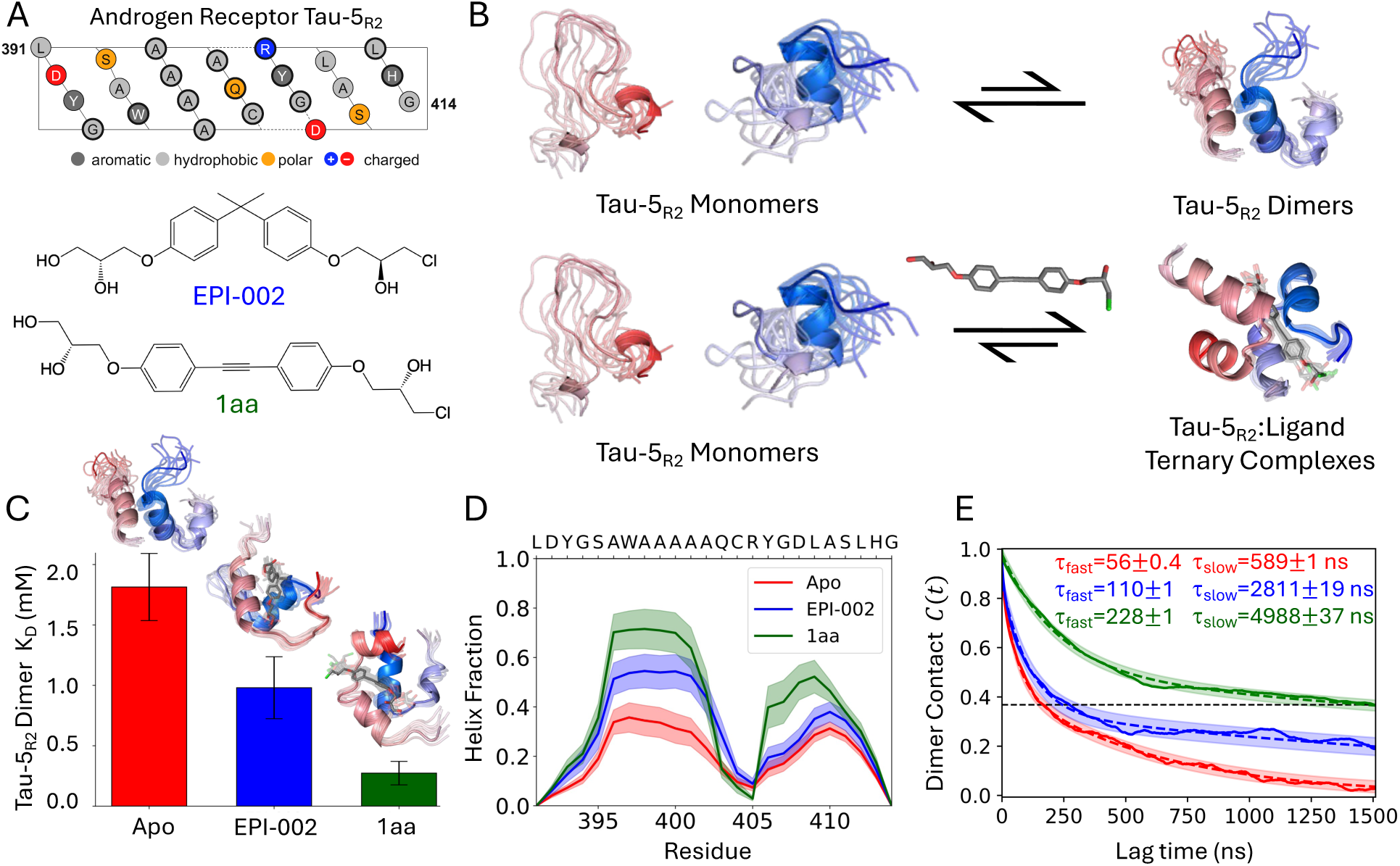
Molecular dynamics simulations of the intermolecular association of Tau-5_R2_ monomers in the presence of EPI-002 and 1aa. (**A**) Amino acid composition of Tau-5_R2_ and chemical structures of EPI-002 and 1aa. **(B)** The presence of ligands drives the equilibrium of Tau-5_R2_ towards the formation of dynamic and heterogeneous dimers. **(C)** Tau-5_R2_ dimer dissociation constants (*K*_D_) observed in unbiased 90–100 *µ*s MD simulations. **(D)** Helical propensities observed in MD simulations of Tau-5_R2_ dimerization. **(E)** Normalized autocorrelation functions (*C*(*t*)) of a binary Tau-5_R2_ dimerization state, where frames containing at least one intermolecular C*α*–C*α* distance below 10Å are assigned a value of 1 (dimer) and all other frames a value of 0 (unbound monomers). These functions quantify how quickly the system loses memory of whether the monomers were associated or dissociated at an earlier time, rather than measuring the continuous lifetime of individual dimers. Double-exponential fits (dashed lines) yield fast (*τ*_fast_) and slow (*τ*_slow_) correlation times.

To elucidate the molecular mechanisms by which AR AD inhibitors stabilize oligomers of the AR AD, we performed all-atom, explicit-solvent MD simulations of the reversible association and dissociation of two Tau-5_R2_ monomers (residues 391–414), a 24-residue segment containing the dominant oligomerization interface of Tau-5^26,27^, in the presence and absence of ligands. Previous biophysical studies of AR AD oligomerization and ligand binding were carried out on Tau-5* (residues 330–448), a disordered 119-residue fragment of the AR AD containing the binding site of EPI-001^5,12,26,27^. Chemical shift perturbations, intermolecular paramagnetic relaxation enhancements, and mutational analyses have identified the Tau-5_R2_ region as the dominant interface mediating Tau-5* oligomerization. Our two-chain Tau-5_R2_ simulations are therefore not intended to reproduce the conformational ensembles of higher-order AR AD oligomers, which dynamic light scattering measurements suggest may contain ten or more Tau-5* monomers^26^. Instead, our two-chain simulations isolate the elementary association of the primary Tau-5 intermolecular oligomerization interface and enable a statistically robust characterization of the kinetics and thermodynamics of Tau-5_R2_ self-association in the presence and absence of AR AD inhibitors at atomic resolution. Simulating two Tau-5_R2_ monomers enables us to observe hundreds-to-thousands of association and dissociation events in 90–100 *µ*s of unbiased all-atom MD simulations. Obtaining comparably extensive sampling of reversible intermolecular association and dissociation of multiple chains of Tau-5* using unbiased all-atom MD is presently infeasible with available computational hardware and sampling methods.

To determine how AR AD ligands alter the kinetics and thermodynamics of Tau-5_R2_ oligomerization, we performed unbiased long-timescale (90–100 *µ*s) all-atom MD simulations in which two Tau-5_R2_ chains reversibly associate and dissociate in the absence of ligand (apo) or in the presence of one molecule of EPI-002 or 1aa (**Figure 1A–B**). We chose to simulate 1aa as its interaction with Tau-5* in solution was previously shown to be stronger than EPI-001 by NMR^12^, while the ligand 1ae was not sufficiently soluble for NMR measurements. We quantified the partitioning of 1aa and EPI-001 into Tau-5* condensates by HPLC, and observed that 1aa has an *∼*3-fold larger partition coefficient (**Supporting Figure S1**), demonstrating stronger enrichment of 1aa in Tau-5* condensates.

Simulations were performed with the a99SB-*disp* protein force field and water model on the Anton 2 supercomputer^28,29^ (“Molecular Dynamics Simulations” in Supporting Information). In each simulation, we classified the two monomers as dimerized when at least one intermolecular C*α*–C*α* distance was below 10Å. The principal conclusions were insensitive to the choice of intermolecular contact distance cutoff. We observed that ligand binding shifts the equilibrium toward associated Tau-5_R2_ states: the simulated *K*_D_ decreases from 1.8 *±* 0.3 mM in the apo simulation to 1.0 *±* 0.3 mM with EPI-002 and 0.30 *±* 0.03 mM with 1aa (**Figure 1C**). Ligand binding also increases the helical population of the associated chains (**Figure 1D**), consistent with experimental observations of increased helical structure in Tau-5_R2_ upon oligomerization^26^.

The trajectories reveal a broad distribution of dissociation timescales for dimers and ternary complexes, rather than the narrower distribution of lifetimes expected for single, well-defined dimer or ternary complex conformational states. Short-lived encounters persist for approximately 1–100 ns, while metastable complexes persist for microseconds, with the longest continuous dimerization event reaching approximately 38 *µ*s in the 1aa simulation (Supporting Figure S2). To quantify the kinetics of Tau-5_R2_ self-association, we calculated autocorrelation functions (ACFs) of the binary dimerization state (**Figure 1E, Supporting Table S1**). These functions quantify how quickly the system loses memory of whether the monomers were associated or dissociated at an earlier time, rather than measuring the continuous lifetime of individual dimers. The ACFs were well described by double-exponential decays with fast (*τ*_fast_ = 50–250 ns) and slow (*τ*_slow_ = 0.5–5.0 *µ*s) relaxation components (**Supporting Table S1**). Both EPI-002 and 1aa increase these correlation times, with 1aa producing the largest increases.

To directly characterize dimer dissociation and reassociation, we additionally calculated the distributions of continuous bound and unbound event durations (**Supporting Figure S2**). Both ligands increase the survival times of Tau-5_R2_ dimers, with 1aa producing a particularly pronounced extension of the long-lived tail of the survival distribution. In contrast, after correcting for differences in the effective protein concentrations of the simulations, the reassociation time distributions are nearly identical for all three systems, suggesting that ligand binding does not substantially alter the kinetics of intermolecular association. Together, these results suggest that EPI-002 and 1aa stabilize Tau-5_R2_ dimers by slowing their dissociation rather than accelerating their association, with 1aa producing the strongest stabilization. As the simulations contain only two Tau-5_R2_ chains at finite concentration, we do not directly interpret these correlation times as experimentally accessible association and dissociation rate constants of Tau-5, and instead interpret the large changes in Tau-5_R2_ interface stability as a measure of the relative effect of ligand binding on Tau-5_R2_ oligomer stability. These simulations suggest that EPI-002 and 1aa enhance Tau-5_R2_ self-association by stabilizing dynamic multivalent oligomerization interfaces, providing a molecular basis by which these ligands and related compounds lower the *T*_C_ of AR condensation and rigidify AR condensates^12,26^.

We compared the conformational properties of Tau-5_R2_ dimer and ternary complexes to identify molecular interactions that distinguish apo and ligand-stabilized complexes (**Figure 2**). To determine how ligand binding alters intermolecular interactions, we calculated residue-resolved intermolecular contact probabilities for the apo, EPI-002, and 1aa simulations (**Figure 2A**). In the apo ensemble, intermolecular contacts are concentrated within the ^393^YGSAWAAAAAQ^403^ region. The addition of EPI-002 increases the helical content and intermolecular contact populations of residues ^396^AWAAAAA^402^, stabilizes aromatic-aromatic interactions throughout the sequence, and introduces weak interactions between C-terminal residues. The effects of 1aa are more pronounced, further increasing the helical content and intermolecular contact populations of the ^396^AWAAAAA^402^ region, while stabilizing stronger interactions with the C-terminal residues that are largely absent in the apo ensemble and only weakly populated in the presence of EPI-002.

**Figure 2:**
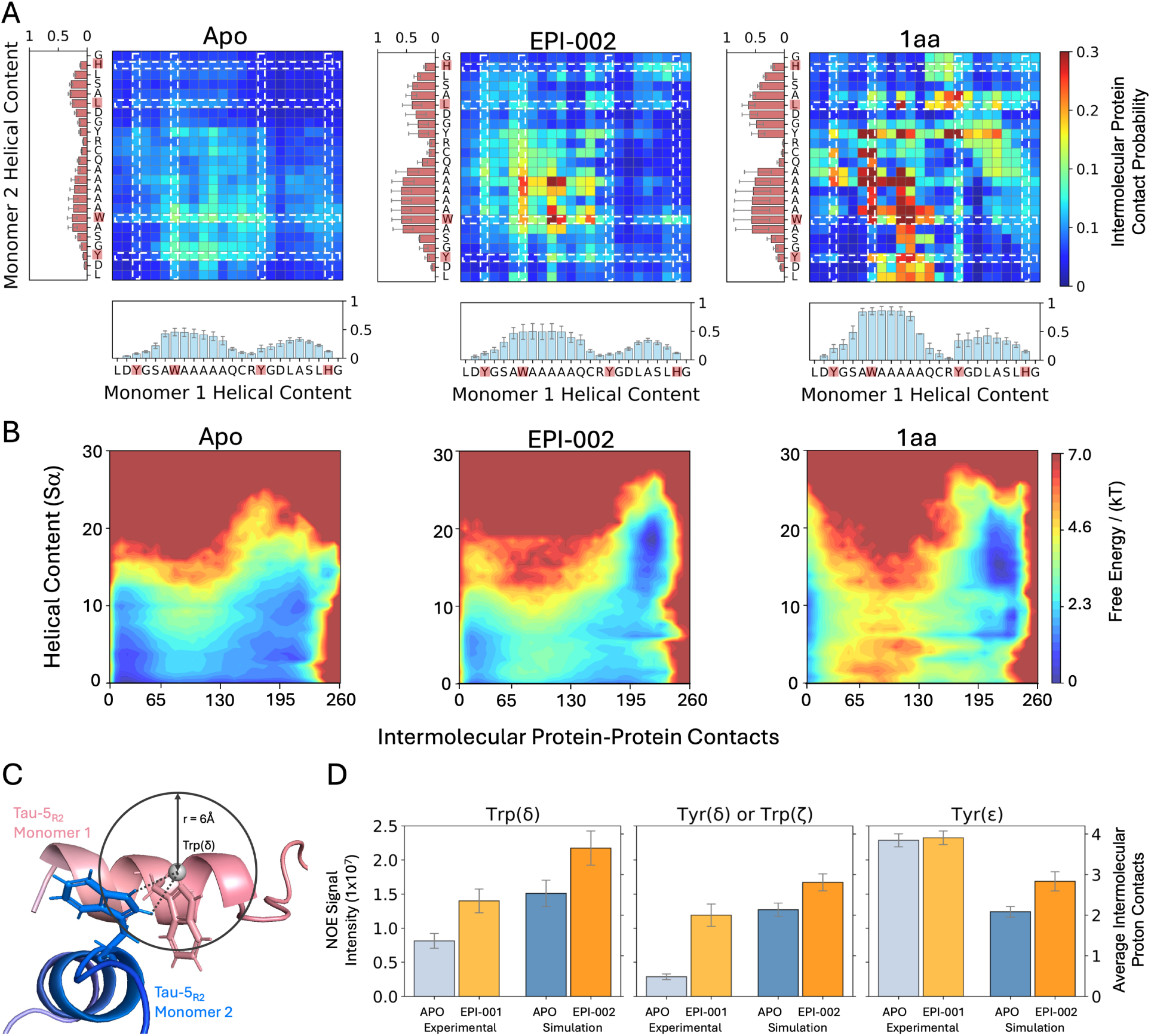
Ligand binding stabilizes aromatic contacts between Tau-5_R2_ monomers. (**A**) Populations of intermolecular contacts between Tau-5_R2_ monomers in unbiased long-timescale MD simulations in both the presence and absence of ligands. Interactions of aromatic residues are highlighted with dashed white lines. Average helical propensities are shown for each monomer. **(B)** Free energy surfaces of Tau-5_R2_ dimerization in each MD simulation as a function of the total *α*-helical content (S*α*) and the number of intermolecular protein–protein contacts, calculated with a smoothed switching function. **(C)** Schematic illustration of intermolecular contacts between an NMR-detected aromatic sidechain proton of a ^13^C-labeled Tau-5* monomer (red) and protons of an unlabeled ^12^C Tau-5* monomer (blue). **(D)** Comparison of experimental intermolecular nuclear Overhauser effect (NOE) intensities measured for aromatic side-chain protons of Tau-5* with the average number of intermolecular proton contacts observed in MD simulations of Tau-5_R2_ dimerization.

We compared the distribution of helical content observed in dimer and ternary Tau-5_R2_ complexes using the helical order parameter *S_α_*, which measures the number of six-residue fragments in a structure that resemble an ideal *α*-helix^30^ (**Figure 2B**). Both ligands increase the population of conformations in which multiple helical segments are formed simultaneously, indicating that ligand binding shifts the ensemble toward states with increased helical structure in both monomers. Notably, 1aa does not substantially increase the maximum helical content relative to EPI-002, but instead strongly reduces the population of non-helical states and increases the population of highly helical conformations. The increased helicity observed with 1aa reflects a redistribution of the ensemble towards more helical states rather than the formation of substantially longer helices. A complex containing two ideal 24-residue *α*-helices would have an *S_α_* value of 38, indicating that even the most helical conformations observed in our simulations remain only partially helical.

The residue-level features of the simulated ensembles reported here are supported by several independent comparisons with previously reported NMR experiments (”Comparing NMR intermolecular chemical shift measurements with molecular dynamics simulations” in Supporting Information). MD-predicted protein-induced chemical shift perturbations reproduce the locations of the major experimental perturbations across Tau-5_R2_ upon oligomerization in the absence and presence of ligand (**Supporting Figure S3**). The distribution of EPI-002 contacts observed in simulation closely resembles the experimental EPI-001-induced chemical shift perturbation profile, supporting the location of the ligand-binding interface identified in the simulations (**Supporting Figure S4**). Additionally, the simulations predict that oligomerization increases *α*-helical propensity primarily within the central Tau-5_R2_ region, in agreement with the residue-resolved increase in *α*-helical propensity observed experimentally in a covalent dimer of two Tau-5* monomers, as inferred from secondary NMR chemical shifts (**Supporting Figure S5**). Together, these comparisons support the location of the simulated protein–protein and protein–ligand interaction interfaces and the accompanying oligomerization-induced increase in local *α*-helical structure.

To experimentally characterize the populations of intermolecular aromatic contacts upon ligand binding, we measured residue-type-specific intermolecular ^13^C-filtered/^13^C-edited nuclear Overhauser effects (NOEs)^31^ for aromatic side-chain protons of Tau-5* in the presence and absence of EPI-001 (**Figure 2C–D**; Supporting Figures S6, S7 and S8; “NMR Spectroscopy” in Supporting Information). In this experiment, isotope filtering suppresses intramolecular signals from the ^13^C-labeled protein while isotope editing selects NOEs originating from ^13^C-bound aromatic protons, allowing NOEs arising from contacts between labeled and unlabeled Tau-5* monomers to be selectively detected. Control experiments showed that the ^13^C filter suppresses *∼*97% of the signal from ^13^C-bound protons and that, in 1:1 ^13^C:^12^C mixtures, NOE cross-peak intensities increase with total protein concentration, confirming that the measured NOEs report on intermolecular contacts (Supporting Figure S8). We observe that the addition of EPI-001 increases several NOE signal intensities resulting from intermolecular contacts between aromatic protons in ^13^C-labeled Tau-5* monomers and unlabeled Tau-5* monomers. These NOE experiments support the ligand-induced oligomerization mechanisms observed in our MD simulations, where EPI-002 and 1aa substantially increase the populations of intermolecular aromatic contacts in a heterogeneous ensemble of intermolecular interfaces.

We compared the relative increase in intermolecular aromatic proton contacts observed upon addition of EPI-002 to our simplified Tau-5_R2_ system with the changes in NOE intensities measured in Tau-5* upon addition of EPI-001 (**Figure 2C–D**, “Comparing NMR intermolecular NOE measurements with molecular dynamics simulations” in Supporting Information) and observe qualitative agreement for several aromatic signals. We note that the experimental Tyr *ε* signal changes little despite an increase in the simulated contacts. We use the MD proton-contact counts as a qualitative structural metric rather than a quantitative back-calculation of NOE intensities, which would require *r^−^*^6^ weighting and explicit modeling of molecular rotational dynamics. Unlike globular proteins, IDPs exhibit heterogeneous, conformation-dependent segmental motions that cannot be described by a single rotational correlation time^32,33^. These dynamics are further complicated by transient oligomerization, and the effective correlation times governing intermolecular NOE cross-relaxation in Tau-5* oligomers are not experimentally established. The experimental NOE measurements were performed using the 119-residue Tau-5* construct, whereas the simulations contain the 24-residue Tau-5_R2_ fragment; differences in construct size and dynamics further preclude a direct quantitative comparison of simulated contact populations with experimental NOE intensities. Due to the limited solubility of 1aa and its effect on Tau-5* saturation concentration, equivalent intermolecular NOE measurements could not be obtained for 1aa.

In addition to consistency with experimental intermolecular aromatic proton NOE measurements and chemical shift perturbations observed upon Tau-5 oligomerization in the presence and absence of EPI-001 (**Supporting Figures S3–S5**), the simulated Tau-5_R2_ oligomerization mechanisms mediated by aromatic contacts and partially folded helical conformations observed here are highly consistent with previously reported NMR measurements and phase separation assays on an extensive library of Tau-5 mutants^12,26^ (**Supporting Tables S2 and S3**, “Consistency with previously reported AR mutational analyses” in Supporting Information). These mutational analyses demonstrated that Tau-5 constructs with diminished helical content and reduced aromatic content have a lower propensity to form oligomers and phase separate in the presence and absence of EPI-001, and substantially reduce the propensity of EPI-001 to partition into Tau-5 condensates.

We next asked whether the long-lived ternary complexes observed in our simulations contain a small number of well-defined ligand-stabilized interfaces or a heterogeneous ensemble containing several distinct interfaces. We described the conformational dynamics of dimer association in each MD trajectory using writhe^34–36^, a geometric descriptor that reports on the relative positions and orientations of protein backbone segments and is well-suited to describe both intramolecular and intermolecular IDP chain organization (“Kinetic Modeling” in Supporting Information). We previously demonstrated that writhe descriptors calculated from protein backbones capture IDP chain organization and provide a substantially more robust description of conformational kinetics than interatomic distances, which are insensitive to the orientation of chain crossings^35^. We performed time-lagged canonical correlation analysis (tCCA) on intramolecular and intermolecular writhe features computed for dimer simulation trajectory data to identify low-dimensional writhe reaction coordinates that describe the slowest dynamical intermolecular association processes observed in each MD simulation. Free energy surfaces of the conformational states sampled in each simulation are shown as a function of the two slowest dynamic modes obtained from tCCA in **Supporting Figures S9, S15 and S22**.

We performed clustering to identify metastable conformational states and constructed Markov state models (MSMs) to quantify the kinetics of transitions between these states in each MD simulation. We display the MSM transition network of Tau-5_R2_ association and dissociation in the presence of EPI-002 in **Figure 3**, with the corresponding networks from the apo and 1aa simulations shown in **Supporting Figures S10 and S23**. The estimated MSMs resolve multiple metastable states connected by exchange on sub-microsecond to multi-microsecond timescales. State-averaged contact maps, helicity, ligand contacts, and chain crossings quantified by writhe are shown in **Supporting Figures S11, S12, S16, S17, S18, S24, S25 and S26**. The properties of each metastable state are summarized in **Supporting Tables S4–S6**. These analyses demonstrate that neither ligand selects for a single dominant dimer structure. Instead, EPI-002 and 1aa stabilize heterogeneous ensembles of interconverting intermolecular interfaces with greater helical content and more extensive networks of intermolecular contacts than apo dimers. MSM implied-timescale and Chapman–Kolmogorov validation tests are provided in **Supporting Figures S13, S20 and S28**, with state-to-state mean first-passage times and transition fluxes shown in **Supporting Figures S14, S21 and S29**. We observe that the presence of ligands substantially slows the rates of interconversion among metastable conformational states relative to the apo simulation.

**Figure 3:**
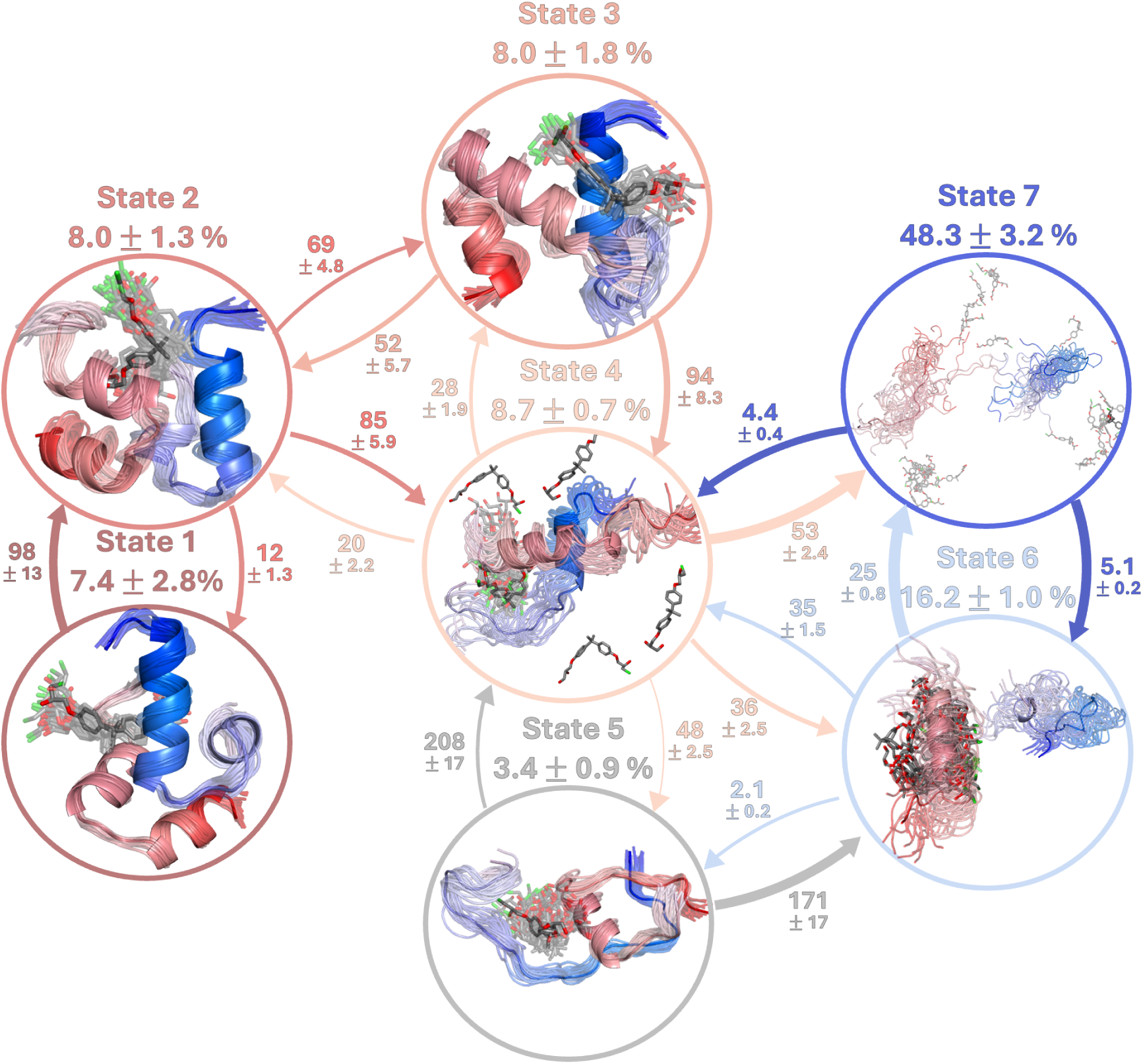
Markov state model (MSM) of Tau-5_R2_ self-association in the presence of EPI-002 estimated from an unbiased 100 *µ*s molecular dynamics simulation. Transition probability fluxes between states are indicated with directed arrows with thickness proportional to the magnitude of the flux. Arrows are labeled with mean first passage times between states, reported in *µ*s. Representative structures of each conformational state are displayed in circles with their steady-state probabilities above. In representative structures of each state, the Tau-5_R2_ monomers are colored with a blue-to-red gradient from the N-terminus of the first monomer to the C-terminus of the second. Error bars indicate 95% bootstrap confidence intervals calculated from 1,000 resamples.

Finally, we asked why 1aa stabilizes Tau-5_R2_ association more strongly than EPI-002. Both ligands preferentially contact aromatic residues, but 1aa forms substantially higher populations of hydrophobic and aromatic-stacking interactions with Tau-5_R2_ (**Figure 4B**; **Supporting Figures S19 and S27**). We quantified “dual residue contacts,” in which a ligand simultaneously contacts residues from opposing Tau-5_R2_ chains. EPI-002 preferentially bridges interfaces involving the ^396^AWAAAAAQ^403^ region (**Figure 4A**), consistent with previously reported NMR measurements^5,12,26^. In contrast, 1aa forms a broader and more highly populated set of bridging interactions. The extended diphenylacetylene linker of 1aa increases the distance between phenyl rings and provides greater rotational flexibility relative to the gem-dimethyl linker of EPI-002. This allows 1aa to more effectively intercalate between the Tau-5_R2_ chains and simultaneously stack with aromatic residues from both monomers, more effectively stabilizing intermolecular interfaces (**Figure 4C**).

**Figure 4:**
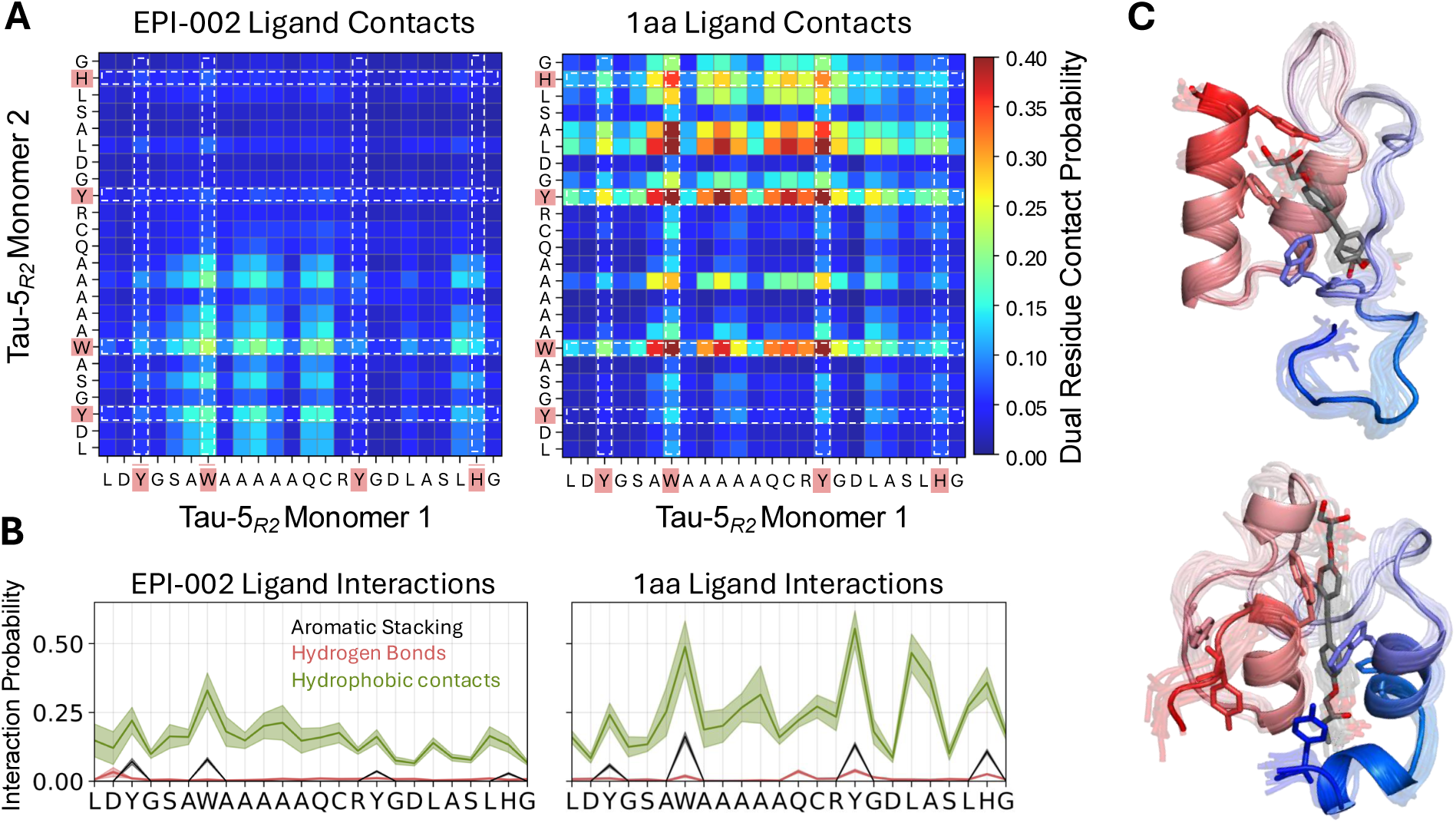
Populations of protein–ligand interactions in molecular dynamics simulations of the intermolecular association of Tau-5_R2_ monomers in the presence of EPI-002 and 1aa. (**A**) Probability that ligands simultaneously form contacts with pairs of residues from opposing Tau-5_R2_ monomers. **(B)** Average populations of protein-ligand interactions between Tau-5_R2_ and either EPI-002 or 1aa. Populations are reported as mean values *±* statistical error estimates from block averaging. **(C)** Representative structures of 1aa forming ternary complexes with two Tau-5_R2_ monomers.

Together, the MD simulations and NMR measurements reported here support a model in which AR AD inhibitors do not stabilize a single rigid Tau-5 oligomerization interface. Instead, they shift a heterogeneous ensemble of dynamic intermolecular interfaces toward longer-lived interfaces with greater helical content, more extensive networks of intermolecular contacts and increased structural order. The mechanisms of ternary complex stabilization reported here identify ligand features and binding modes that stabilize AR AD oligomers and provide an atomic-level explanation for the increased potency of 1aa and other diphenylacetylene derivatives in AR transcriptional inhibition assays compared to EPI-001^12^. Because the simulations isolate one recurring Tau-5 interface rather than the complete higher-order assembly, we interpret these structures as elementary building blocks that can be reused within larger Tau-5 oligomers and condensates.

Recent work has shown that EPI-001 binds the AR AD with greater affinity than four other steroid-receptor ADs and selectively stabilizes AR AD oligomers without stabilizing oligomers of the other ADs^26^. This suggests that, in some systems, ligand recognition of oligomeric interfaces—rather than monomeric states—may be a major determinant of IDP ligand selectivity. In such cases, designing ligands that preferentially bind and stabilize specific oligomeric conformations could provide a general strategy for enhancing IDP ligand affinity and selectivity. Mechanistic insight from all-atom MD simulations of ligand-mediated IDP oligomerization, as provided here, may prove valuable for guiding this emerging structure-based drug design paradigm for IDPs.

Lastly, we speculate that the mechanisms of ligand-mediated oligomerization of Tau-5_R2_ reported here, together with interactions involving additional segments of Tau-5, may underlie the increased rigidity of AR condensates observed in the presence of EPI-001^26^. In future studies, we will investigate this hypothesis by attempting to leverage mechanistic insights from all-atom MD simulations of ligand-mediated IDP oligomerization to design small molecules that rationally tune the material properties of biomolecular condensates. In summary, the atomic-resolution mechanistic framework presented here will inform ongoing efforts to develop therapeutics for the treatment of CRPC and other currently untreatable diseases associated with IDP and condensate dysfunction^37,38^.

## Data and Code Availability

All molecular dynamics simulation input files and analysis code are freely available from https://github.com/paulrobustelli/Zhu_Sisk_AR_oligomers_2025. The MD trajectories are publicly available through the data archive linked in the repository.

## Acknowledgments

P.R., J.Z. and T.R.S. acknowledge the support of NIGMS award R35GM142750. T.R.S. additionally acknowledges the support of the U.S. Department of Education award GAANN P200A240037. Anton 2 computer time was provided by the Pittsburgh Supercomputing Center (PSC) through Grant R01GM116961 from the National Institutes of Health and Anton award MCB200087P. The Anton 2 machine at PSC was made available by D.E. Shaw Research.

B.M. acknowledges the support of the AECC (Spanish Cancer Association) postdoctoral grant. S.B.G. acknowledges support from a PhD fellowship awarded by IRB in the 2020 call and an FPI fellowship awarded by MINECO in the 2021 call (PID2019-110198RV-100). X.S. was supported by AGAUR (2021 SGR 476), the AEI (PID2019-110198RB-I00 and PID2022-141816OB-I00), the Fundació La Caixa (CI20-00098), the AECC (INNO20010FRIG) and the Mark Foundation.

