## Supporting Information for "How small molecules stabilize oligomers of a phase-separating disordered protein"

Jiaqi Zhu,[†,§] Thomas R. Sisk,[†,§] Borja Mateos,[‡] Stasė Bielskutė-Garcia,[‡] Xavier Salvatella,[‡,¶] and Paul Robustelli[*,†]

†*Department of Chemistry, Dartmouth College, Hanover, NH, 03755, USA*

‡*Institute for Research in Biomedicine (IRB Barcelona), The Barcelona Institute of Science and Technology, Baldiri Reixac 10, 08028 Barcelona, Spain*

¶*ICREA, Barcelona, Spain*

§*These authors contributed equally to this work.*

### Supporting Information

#### Molecular dynamics simulations.

Initial structures of Tau-$5_{R2}$ (residues L391-G414, capped with ACE and NH2 groups at the N and C termini, respectively) were generated using the pmx software package[1]. The builder module of pmx was used to generate a fully helical Tau-$5_{R2}$ conformation, where all residues were set in an ideal helical structure ($\phi = -57°, \psi = -47°$). These structures were subjected to an energy minimization step, followed by a short 100 ps high temperature unfolding simulation at 600 K in the NVT ensemble to generate a diverse set of unfolded conformations. A final structure was selected from the unfolding trajectories for further simulation.

The selected structure underwent energy minimization using the steepest descent algorithm until the maximum force was reduced to less than 1000.0 kJ/(mol·nm). Equilibration was performed in two stages: first, a 2000 ps NVT equilibration at 300 K using the Berendsen thermostat[2], followed by a 200 ps NPT equilibration at 1 bar with the Martyna-Tobias-Klein barostat while maintaining the temperature at 300 K. Position restraints were applied to all heavy atoms to allow solvent relaxation while preserving protein structure. Bond lengths and angles of protein atoms were constrained using the LINCS algorithm[3], while water constraints were enforced using the SETTLE algorithm[4]. Long-range electrostatic interactions were calculated using the Particle Mesh Ewald (PME) algorithm[5] with a grid spacing of 0.16 nm, and van der Waals interactions were truncated at 0.9 nm.

All molecular dynamics simulations were performed using an all-atom explicit solvent model. The Tau-$5_{R2}$ dimer was prepared in GROMACS 2022[6,7], starting from a single monomer that was energy minimized and subsequently duplicated using the gmx insert-molecules command to generate the dimer. We employed the a99SB-*disp* protein force field[8], a99SB-*disp* water model[8] and CHARMM22* ion parameters[9]. Each system was solvated in a cubic box with a99SB-*disp* water: the apo Tau-$5_{R2}$ dimer was solvated in an

80×80×80 Å$^3$ box containing 16,386 water molecules, the EPI-002 system in an 80×80×80 Å$^3$ box containing 16,356 water molecules, and the 1aa system in a 110 × 110 × 110 Å$^3$ box containing 42,889 water molecules. All systems were neutralized with CHARMM22* ions[9], and additional $Na^+$ and $Cl^-$ ions were added to achieve a final bulk salt concentration of 20 mM NaCl prior to equilibration and production simulations. Ligands were parameterized using the GAFF1 forcefield[10].

To enable simulations on the Anton 2 supercomputer, the solvated system was converted from GROMACS format to Desmond format using InterMol[11]. The generated CMS file was then processed using the VIPARR software to apply the appropriate force field parameters. Once each system was fully assembled, a short equilibration run was performed using an integration time step of $dt = 0.0025$ ps. After equilibration, hydrogen mass repartitioning[12] was applied. Hydrogen atoms were assigned a mass of 4 amu and oxygen atoms in water molecules were assigned a mass of 10 amu to enable the use of 4 fs integration time steps. All systems used the same mass-repartitioning and integration protocol. We therefore interpret kinetic differences primarily as relative differences among the simulated conditions rather than as experimentally exact absolute timescales. The finite simulation volumes set the effective protein concentrations and influence bimolecular encounter frequencies; accordingly, simulated association times are not interpreted as experimental bimolecular association rate constants.

Production simulations were conducted on the Anton 2 supercomputer[13] at the Pittsburgh Supercomputing Center (PSC). Long-range nonbonded interactions were updated using the RESPA multiple time-step integration scheme. Simulations of two Tau-$5_{R2}$ monomers in the absence of ligands (apo) or in the presence of one molecule of EPI-002 were performed for 100 $\mu$s each. The corresponding simulation containing one molecule of 1aa was performed for 90 $\mu$s due to Anton 2 allocation limits. Analyses were run utilizing MDtraj[14] and the numpy[15] python package.

#### Molecular dynamics simulation analyses.

The stability of the Tau-$5_{\mathrm{R2}}$ dimer was quantified by analyzing intermolecular C$\alpha$ distances between Tau-$5_{\mathrm{R2}}$ monomers in the simulations. We classified the two monomers as dimerized when at least one intermolecular C$\alpha$–C$\alpha$ distance was below 10 Å. For free energy surfaces in Figure 2B, we utilize a smoothed quantification of intermolecular contacts by passing pairwise intermolecular C$\alpha$ distances through a sigmoid switching function centered at 10 Å before summing over all residue pairs, $\sum_{i,j} 1 - \frac{1}{1+e^{-(d_{i,j}-1)}}$. In the same figure, we obtain a smoothly varying estimate of the $\alpha$-helical content in the Tau-$5_{\mathrm{R2}}$ monomers via the order parameter S$\alpha$[16], which is computed using the root mean square deviation ($RMSD_{\alpha_i}$) between each seven-residue fragment of simulated conformations and an ideal $\alpha$-helix composed of the same residues. Each seven-residue fragment is assigned a value between 1 (perfect $\alpha$-helix) and 0 (not $\alpha$-helical) by passing the RMSD through the switching function:

$$S_{\alpha_i} = \frac{1 - \left(\frac{\mathrm{RMSD}_{\alpha_i}}{r_0}\right)^8}{1 - \left(\frac{\mathrm{RMSD}_{\alpha_i}}{r_0}\right)^{12}} \tag{S1}$$

The threshold parameter, $r_0$, is set to 1.0 Å such that fragments with RMSDs from an ideal $\alpha$-helix beyond this rapidly decay to zero and therefore do not contribute to the estimated $\alpha$-helical content. In addition, we utilize the DSSP algorithm[17] to obtain a binary classification of individual residues as being in an $\alpha$-helical conformation (1) or not (0). The ensemble averaged $\alpha$-helical fraction obtained using the DSSP algorithm for each MD simulation is reported in Figure 1D.

Dimer dissociation $K_D$ values were calculated according to $K_D/c^\circ = (P_u/P_b)(\nu c^\circ N_A)^{-1}$, where $P_u$ and $P_b$ are the fractions of frames classified as unbound and dimeric, respectively, $\nu$ is the simulation box volume, $N_A$ is Avogadro's number, and $c^\circ$ is the standard state concentration (1 mol $\mathrm{L}^{-1}$). The simulated concentration of two copies of Tau-$5_{\mathrm{R2}}$ is 6.66 mM in the 80 Å box used for the apo and EPI-002 simulations and 2.56 mM in the 110 Å box used for the 1aa simulation.

Dimer binding events were identified from consecutive saved frames containing at least one intermolecular C$\alpha$–C$\alpha$ minimum-image distance below 10 Å. The distributions of observed binding-event durations, dimer survival, and concentration-scaled reassociation times are shown in Supporting Figure S2 and defined below. The final 1aa dimer remained associated for at least 37.9 $\mu$s; dissociation was not observed before the trajectory ended. The longest completed binding events in the apo and EPI-002 simulations lasted 5.3 and 15.1 $\mu$s, respectively.

We defined ternary complexes as occurring in frames in which both Tau-5$_{\mathrm{R2}}$ monomers are simultaneously in contact with a ligand. We defined Tau-5$_{\mathrm{R2}}$ monomers to be in contact with a ligand if any protein heavy atom was within 6 Å of a ligand heavy atom. The lifetimes of ternary complex were determined by identifying consecutive frames containing ternary complexes. We observe that the most stable ternary complex formed with 1aa has a lifetime of 28.7 $\mu$s. This ternary complex represents a continuous subset of the frames comprising the 37.9 $\mu$s Tau-5$_{\mathrm{R2}}$ dimerization event observed in this simulation. The most stable ternary complex formed with EPI-002 has a lifetime of 9.5 $\mu$s. Similarly, this ternary complex represents a continuous subset of the frames from the most stable 15.1 $\mu$s Tau-5$_{\mathrm{R2}}$ dimerization event from this simulation.

Protein-ligand contacts were defined as occurring in any frame where at least one heavy (non-hydrogen) atom of a residue is found within 6.0 Å of a ligand-heavy atom. Protein-ligand hydrophobic contacts were defined as occurring when pairs of protein and ligand carbon atoms were within 4 Å. Potential hydrogen bond donors were defined as nitrogen, oxygen, or sulfur atoms bearing an attached hydrogen; potential acceptors were defined as nitrogen, oxygen, or sulfur atoms. Hydrogen bonds were identified with a distance cutoff of 3.5 Å between the donor-hydrogen and heavy-atom acceptor, and a donor-hydrogen-acceptor angle $> 150°$.

Aromatic stacking interactions between ligands and protein aromatic side chains were calculated as previously described[18]. For a protein aromatic ring and a ligand aromatic ring: we define $R$ as the distance between ring centers, $\hat{R}$ as the unit vector connecting the ring centers, $\hat{n}_{\text{protein}}$ and $\hat{n}_{\text{ligand}}$ are normal vectors to the side chain and ligand ring planes originating from the ring centers, $\theta$ is the angle between $\hat{n}_{\text{protein}}$ and $\hat{n}_{\text{ligand}}$ ($\theta = \arccos(\hat{n}_{\text{protein}} \cdot \hat{n}_{\text{ligand}})$), and $\varphi$ is the angle between $\hat{n}_{\text{protein}}$ and $\hat{R}$ ($\varphi = \arccos(\hat{n}_{\text{protein}} \cdot \hat{R})$). Parallel stacked conformations were defined as occurring when $R < 6.5$ Å, $\theta < 45°$, and $\varphi < 60°$ and T-stacked conformations were defined as occurring when $R < 7.5$ Å, $\theta > 75°$, and $\varphi < 60°$.

We quantified the probability of simultaneously forming ligand contacts in all pairs of residues from opposing Tau-$5_{\text{R2}}$ monomers (Figure 4, Supporting Figures S17 and S25) and refer to this quantity as the "dual residue contact probability".

##### Double-exponential fitting of dimer-state autocorrelation functions

To characterize the timescales of Tau-$5_{\text{R2}}$ association/dissociation exchange, we calculated normalized autocorrelation functions from binary time series describing the presence or absence of intermolecular protein contacts. The mean was subtracted from each binary time series before calculating the autocorrelation function, which was normalized by the zero-lag variance. These functions quantify association-state correlations and allow intervening dissociation and reassociation; the fitted times are therefore correlation times rather than continuous dimer lifetimes. The autocorrelation functions were fit to a double-exponential decay,

$$C(t) = a_{\text{fast}} \exp\left(-\frac{t}{\tau_{\text{fast}}}\right) + a_{\text{slow}} \exp\left(-\frac{t}{\tau_{\text{slow}}}\right), \tag{S2}$$

where $a_{\text{fast}}$ and $a_{\text{slow}}$ are the fitted amplitudes and $\tau_{\text{fast}}$ and $\tau_{\text{slow}}$ are the corresponding correlation times. All amplitudes and correlation times were constrained to be non-negative during fitting. The fitted parameters for the apo, EPI-002, and 1aa simulations are reported in Supporting Table S1.

**Supporting Table S1.** Double-exponential fit parameters for Tau-$5_{\mathrm{R2}}$ dimer-state autocorrelation functions. Reported uncertainties are $\pm 1\sigma$ standard errors obtained from the covariance matrix of the nonlinear least-squares fit. These errors quantify fit precision and do not account for finite-trajectory sampling uncertainty.

| System | $a_{\mathrm{fast}}$ | $\tau_{\mathrm{fast}}$ (ns) | $a_{\mathrm{slow}}$ | $\tau_{\mathrm{slow}}$ ($\mu$s) | $R^2$ |
|---|---|---|---|---|---|
| Apo | $0.4192 \pm 0.0017$ | $56.05 \pm 0.45$ | $0.4580 \pm 0.0011$ | $0.5886 \pm 0.0014$ | 0.9936 |
| EPI-002 | $0.5264 \pm 0.0017$ | $109.85 \pm 0.71$ | $0.3390 \pm 0.0009$ | $2.8111 \pm 0.0192$ | 0.9776 |
| 1aa | $0.4865 \pm 0.0010$ | $228.26 \pm 1.04$ | $0.4930 \pm 0.0010$ | $4.9883 \pm 0.0366$ | 0.9939 |

#### Dimer survival and concentration-scaled reassociation

We identified binding events and reassociation times from trajectories saved every $\Delta t$ = 0.24 ns. A frame was classified as bound when at least one intermolecular C$\alpha$–C$\alpha$ minimum-image distance was below 10 Å, and as unbound otherwise. Binding-event durations were calculated as the number of consecutive bound frames multiplied by $\Delta t$. Reassociation times were calculated in the same way from consecutive unbound frames following dissociation. The initial unbound segment of each trajectory was excluded from the reassociation analysis because no preceding dissociation was observed.

This analysis identified 3,879, 2,441, and 558 binding events in the apo, EPI-002, and 1aa simulations, respectively (Supporting Figure S2A). Dissociation was observed for all events except the final 1aa event, which remained associated for at least 37.927 $\mu$s before the trajectory ended. These counts include brief contacts and rapid recrossings of the distance threshold; they do not represent an equal number of statistically independent samples.

To describe dimer survival, we calculated the fraction of binding events remaining associated as a function of time since association. Events still ongoing at the end of a trajectory were included for their observed duration, without treating the trajectory endpoint as a dissociation. The survival fraction was calculated as

$$S(t) = \prod_{t_j \leq t} \left(1 - \frac{d_j}{n_j}\right),$$

where $d_j$ is the number of dissociations at duration $t_j$ and $n_j$ is the number of events that remained associated and were still being followed immediately before that duration. If a dissociation and a trajectory endpoint occurred at the same event duration, both events were included in $n_j$, but only the dissociation contributed to $d_j$. Supporting Figure S2B shows the initial 10 ns on linear axes; panel C shows the long-lived tail with a logarithmic vertical axis. The unfinished 1aa event is marked with an asterisk.

To compare reassociation across simulation boxes, we scaled unbound durations to the mean apo partner concentration. For two monomers, the concentration of the single available binding partner in frame $k$ is $c_k = 1/(N_\mathrm{A} V_k)$, where $V_k$ is the box volume in litres and $N_\mathrm{A}$ is Avogadro's constant. The scaled duration is

$$t_\mathrm{scaled} = \frac{\Delta t}{\bar{c}_\mathrm{apo}} \sum_{k \in \mathrm{unbound\ event}} c_k,$$

where $\bar{c}_\mathrm{apo}$ is the mean apo partner concentration. At constant volume, this reduces to $t_\mathrm{scaled} = t_\mathrm{unbound} c/\bar{c}_\mathrm{apo}$. Mean partner concentrations were 3.327, 3.331, and 1.280 mM for apo, EPI-002, and 1aa, respectively. The correction therefore leaves EPI-002 waiting times nearly unchanged and scales 1aa waiting times by approximately 0.385. This comparison assumes that association frequency is proportional to partner concentration.

Supporting Figure S2D shows the fraction of events remaining unbound as a function of concentration-scaled time since dissociation. These curves were calculated using the same survival expression, with reassociation replacing dissociation as the event that reduces the surviving fraction. Unbound events ongoing at the trajectory endpoint contributed for their observed, concentration-scaled duration without counting the endpoint as a reassociation.

#### Kinetic Modeling and Markov State Model Estimation

Markov state models were estimated from trajectories with a saved-frame spacing of 0.24 ns. We characterized the conformational states of Tau-$5_\mathrm{R2}$ dimers and ternary complexes by analyzing fluctuations in intramolecular and intermolecular backbone chain organization using the writhe, a geometric measure from knot theory that quantifies signed crossings of three-dimensional (3D) curves[19–21]. Our computation of the writhe yields pairwise scalar descriptors between backbone segments that quantify both their proximity and relative orientations. Unlike an interatomic distance, the signed crossing distinguishes not only whether two backbone segments are spatially close but also their relative orientation. For a two-chain system, these descriptors can be calculated between a backbone segment from one monomer and a segment from the other monomer, so the intermolecular writhe terms describe the relative geometry and orientation of the two chains at the oligomerization interface. We visualize average intermolecular and intramolecular chain crossings in each metastable conformational state of MSMs estimated for the apo, EPI-002 and 1aa dimer simulations in Supporting Figures S12, S18, and S26, respectively. Sisk et al.[21] provide a detailed description of this calculation and demonstrate that writhe descriptors are effective at identifying slow processes in MD simulations of IDPs compared with traditional order parameters such as Euclidean distances or torsion angles. We performed all calculations of the writhe from MD data using the `writhe-tools` Python package which is available on the Python Package Index (PyPI).

To obtain a quantitative description of the kinetic properties of Tau-$5_{\mathrm{R2}}$ dimers and ternary complexes observed in MD simulations, we construct Markov state models (MSMs)[22]. Here, we build MSMs by first reducing the dimension of intramolecular and intermolecular writhe features using time-lagged canonical correlation (tCCA) analysis[21,23], a dimensionality reduction method that finds slowly evolving linear combinations of input features that yield optimal reaction coordinates for MSM construction[22]. In Supporting Figures S9, S15, and S22, we visualize 2D tCCA projections computed for each MD ensemble and color projected MD frames by free energy and additional reaction coordinates including protein-protein contacts and helical content.

To identify MSM states, we applied K-means clustering[24] to the low-dimensional tCCA projections and subsequently estimated MSMs using the `deeptime` Python package.[25] To determine the optimal number of tCCA reaction coordinates and K-means clusters, we performed a grid search over 2–10 tCCA dimensions and 40–500 clusters. For each set of hyper-parameters, we estimated an MSM and computed the corresponding implied time scales (ITS) and VAMP1 score[23]. We selected model hyperparameters based on the highest VAMP1 score and convergence of the implied timescales (ITS), which provide a validation test for MSMs. For the Apo, EPI-002 and 1aa MD simulations, we used 4, 3, and 3 tCCA dimensions, respectively, and 150, 400 and 250 clusters, respectively. We coarse-grained each model to obtain metastable states using the generalized PCCA spectral clustering algorithm[26,27]. The number of metastable states used to coarse-grain an MSM is generally set equal to 1 plus the number of converged ITS where the extra state accounts for the stationary process. Here, we found that the best performing MSMs for all MD simulations exhibited at least 6 converged ITS (Supporting Figures S13, S20 and S28). To enable direct comparisons between MSMs, we used 7 metastable states for all MSMs and a shared lag-time of 120 ns which was the shortest lag-time at which the ITS for all models converged to a Markovian regime. The tCCA decomposition also requires a lag-time parameter. Because we found that this parameter had little effect on the resulting projections, we used a lag time of 150 ns for all systems. We note that due to minimal sampling out of states 1-4 in the 1aa:Tau-$5_{\mathrm{R2}}$ dimer MSM, we found that the MSM predicted steady-state populations deviate from the empirical state counts. Consequently, we estimate the MFPTs and transition probability fluxes from an identical MSM with the steady-state distribution constrained[25] to match empirical populations to ensure reported values are consistent with the original trajectory data.

In Supporting Figure S10, Figure 3 and Supporting Figure S23 we show transition network representations of the MSMs estimated for each system, where representative structures of each metastable state are visualized along with their stationary populations and probabilities are depicted through the width of arrows between states which are labeled with the corresponding mean first passage time (MFPT). Intermolecular protein contact probabilities and $\alpha$-helix propensities in each metastable state of the apo, EPI-002 and 1aa MSMs are shown in Supporting Figures S11, S16, and S24. Protein-ligand contact and the dual-residue contact probabilities in each metastable state of the EPI-002 and 1aa MSMs are shown in Supporting Figures S17 and S25. CK-tests, ITS as a function of the lag-time and steady-state populations obtained from the apo, EPI-002 and 1aa MSMs are shown in Supporting Figures S13, S20, and S28. Transition probability fluxes and MFPTs between metastable states in each model are shown in Supporting Figures S14, S21 and S29. In Supporting Tables S4, S5 and S6 we list the state populations, average number of protein contacts, average number of seven-residue helical segments (S$\alpha$) and population of the dimer complex in each metastable state of the MSMs estimated for each system. For the EPI-002 and 1aa MSMs, we also include the average number of ligand contacts and population of the ternary complex in each metastable state (Supporting Tables S5 and S6).

#### Protein expression and purification

Recombinant expression of Tau-5* was performed in E. coli Rosetta (DE3) cells. Non-isotopically labeled protein was expressed in Lysogeny Broth (LB), whereas isotopically $^{13}$C- or $^{15}$N,$^{13}$C-labeled protein was expressed in minimal M9 media supplemented with $^{13}$C-glucose or with $^{13}$C-glucose and $^{15}NH_4Cl$, respectively. Cell cultures were grown to an $OD_{600}$ of 0.6, induced with 1 mM isopropyl $\beta$-D-1-thiogalactopyranoside (IPTG), and incubated at 25°C overnight under agitation for expression.[28].

Cells were harvested by centrifugation and resuspended in phosphate-buffered saline (PBS). The cells underwent two rounds of sonication for 7 min each with a 5s on/off pulse cycle. Following centrifugation, the supernatants were discarded, and the pellets were washed twice using a wash buffer (PBS, 1% Triton, 500 mM NaCl, 1 mM dithiothreitol (DTT), pH 7.8). For the first wash, 5 mM $MgSO_4$, 130 $\mu$M $CaCl_2$ and 10 µg/mL DNAse I were added. Insoluble inclusion bodies were collected by centrifugation and solubilized overnight at room temperature in binding buffer (20 mM Tris, 500 mM NaCl, 5 mM imidazole, 8 M urea, 0.05% w/v $NaN_3$, 1 mM DTT, pH 7.8). The solubilized material was centrifuged, the supernatant filtered, and applied to a HisTrap HP column (Cytiva) at room temperature. Bound protein was eluted with a gradient to 500 mM imidazole in binding buffer.

Eluted fractions underwent two rounds of dialysis, of 3 h and 16 h, against 50 mM Tris, 1 mM DTT, pH 8.0. His-tag cleavage was performed with 3C protease during the second dialysis. After dialysis, urea was added to a final concentration of 8 M and the sample was applied to a second HisTrap HP column (Cytiva) at room temperature to collect the cleaved protein in the flow-through. Concentration was performed using 3 kDa Amicon® concentrators (Merck). The samples were aliquoted and stored at −80°C. For use, aliquots were thawed and subjected to SEC using HiLoad Superdex 75 pg columns (Cytiva) in 20 mM sodium phosphate, 1 mM tris(2-carboxyethyl)phosphine (TCEP), 0.05% w/v $NaN_3$, pH 7.4, at 4°C.

#### Small molecule partition coefficients

Concentrations of EPI-001 and 1aa in the dense and light phases of Tau-5* were determined using an Agilent Technologies 1200 HPLC instrument, using a Jupiter analytical C4 column from Phenomenex. $H_2O$ and ACN:$H_2O$ (9:1) were used as mobile phases, containing 0.1% TFA. Samples were prepared on ice in a 20 mM sodium phosphate buffer (pH 7.4), 1 mM TCEP, 0.05% $NaN_3$. The detection wavelength was 215 nm.

One molar equivalent of compound was added to 60 $\mu$M protein from DMSO stocks. The final concentration of DMSO in all samples was 2%. EPI-001 or 1aa was first added to the corresponding protein samples, followed by 1.25 M NaCl to induce condensation. This order of addition ensured that the compound was present during condensate nucleation and growth, thereby avoiding potential kinetic barriers to diffusion into preformed condensates and minimizing biases arising from condensate material properties. Samples were incubated for 5 min at 37 °C, followed by centrifugation at 2,000 rpm for 2 min at 37 °C to separate the light and dense phases. The light phase was transferred to a new microcentrifuge tube, while the dense phase was diluted ninefold with buffer containing 4 M urea, which was used to dissolve condensates. Dense- and light-phase samples were injected into the HPLC. The corresponding peaks of the small molecules were integrated, and the ratio of their peak areas was calculated after correcting the dense-phase peak areas for the dilution factor. Measurements were repeated in triplicate.

#### NMR Spectroscopy

Two-dimensional $^{1}$H-$^{13}$C correlation experiments[29] centered at the aromatic region were obtained at 278 K in a Bruker Avance III 600 MHz spectrometer equipped with a TCI cryoprobe using 100–400 $\mu$M $^{15}$N, $^{13}$C-labeled Tau-5* in 20 mM sodium phosphate, pH 7.4, 1 mM TCEP, 0.05% w/v $NaN_3$, 10 $\mu$M sodium 3-(trimethylsilyl)propane-1-sulfonate (DSS). Aromatic side chain $^{1}H_d$ resonances were assigned by acquiring a (Hb)Cb(CgCd)Hd experiment[30] linking directly the $C_b$ to the $H_d$ chemical shifts of aromatic residues, using 100 $\mu$M $^{15}$N,$^{13}$C-labeled Tau-5*. $C_b$ chemical shifts were previously assigned (BMRB ID:53105)[31]. Additional $^{1}$H aromatic resonances were assigned using three-dimensional $^{15}$N-edited NOESY-HSQC and TOCSY-HSQC experiments acquired on 400 $\mu$M $^{15}$N-labeled Tau-5*, together with a 3D $^{13}$C-edited NOESY-TROSY experiment optimized for aromatic residues acquired using 300 $\mu$M $^{15}$N,$^{13}$C-labeled Tau-5* in 200 mM NaCl. Mixing times of 200 and 75 ms were used for the NOESY and TOCSY experiments, respectively. 3D experiments were acquired with non-uniform sampling (NUS) and processed with qMDD[32].

Intermolecular NOEs (with and without EPI-001) were recorded on mixed-state samples containing a total protein concentration of 0.3 mM, composed of $^{13}$C,$^{1}$H-labelled and $^{12}$C,$^{1}$H-labelled Tau-5* in a 1:1 ratio. Samples were prepared in 20 mM sodium phosphate buffer, pH 7.4, containing 1 mM TCEP, with 100% $D_2O$ as solvent. EPI-001 was added from a 50 mM DMSO-$_{d6}$ stock solution to a final concentration of 0.3 mM. Under these experimental conditions, the $C_{sat}$ of the protein construct was 0.7 mM in the absence of EPI-001 and

0.4 mM in its presence. To avoid phase separation and sample instability during NMR measurements, a protein concentration of 0.3 mM (below $C_{sat}$) was used. Lower protein concentrations result in insufficient NOE cross-peaks. Compound 1aa exhibited limited solubility and further decreased the $C_{sat}$ of the protein, rendering the $^{13}$C-filtered/$^{13}$C-edited NOESY experiment technically unfeasible.

NMR NOE experiments were recorded at 800 MHz on a Bruker Avance Neo spectrometer equipped with a TCI cryoprobe at 278 K, using the $^{13}$C-filtered/$^{13}$C-edited pulse sequence of Zwahlen *et al.*[33]. Two-dimensional $^{1}$H–$^{1}$H spectra were acquired with a 200 ms mixing time and a recycle delay $d_1$ of 1.5 s. Samples were measured in 3 mm NMR tubes with 10 $\mu$M DSS. Data were processed using NMRPipe[34] and analyzed with CcpNMR Analysis[35].

#### Controls for the measurement of intermolecular NOEs

Intermolecular NOEs between Tau-5* monomers were measured with the $^{13}$C-filtered/$^{13}$C-edited NOESY pulse scheme of Zwahlen *et al.*[33] In this experiment, magnetization of protons bound to $^{13}$C is removed by a $^{13}$C filter applied before the NOE mixing period, so that only $^{12}$C-bound protons are frequency labeled in the indirect $^{1}$H dimension. After the mixing period, a $^{13}$C-editing element selects magnetization residing on $^{13}$C-bound protons for detection. In mixed samples of $^{13}$C,$^{1}$H- and $^{12}$C,$^{1}$H-labeled Tau-5*, cross-peaks therefore arise from NOEs connecting protons of unlabeled and labeled chains, that is, from intermolecular contacts. The reliability of this approach depends critically on the efficiency of the $^{13}$C filter: any $^{13}$C-bound magnetization that survives the filter gives rise to intramolecular cross-peaks that cannot be distinguished from intermolecular ones. We therefore carried out two sets of control experiments (Supporting Figure S8).

**Efficiency of the $^{13}$C filter.** A double $^{13}$C purge element was used, with purge durations of 4.4 ms ($\tau_a = 2.2$ ms) and 4.0 ms ($\tau_a = 2.0$ ms). Each element employed a WURST adiabatic $^{13}$C inversion pulse centered at 0 ppm and swept over 80 kHz from high to low frequency, of duration 2.358 ms (sweep rate $3.39\times10^7$ Hz/s) and 1.527 ms (sweep rate $5.24\times10^7$ Hz/s), respectively. The adiabatic sweep exploits the approximately linear relationship between one-bond $^1$H–$^{13}$C scalar couplings ($^1J_{\mathrm{CH}}$) and $^{13}$C chemical shifts, so that aliphatic and aromatic $^{13}$C-bound protons are suppressed in a single element.[33] Filter performance was evaluated by comparing one-dimensional $^1$H spectra recorded with and without the $^{13}$C filter on a 0.4 mM sample of uniformly $^{13}$C-labeled Tau-5* (1:0 $^{13}$C:$^{12}$C; Supporting Figure S8A, left). With the filter applied, ~97% of the total $^1$H signal is suppressed. The residual signal (~3%) is attributed mainly to $^{12}$C-bound protons arising from incomplete $^{13}$C incorporation in the labeled protein, and defines the background expected from the labeled component. In the aromatic region, where the intermolecular NOEs analyzed in this work are detected, ~10% of the signal remains after filtering. The largest residual intensity corresponds to His H$\varepsilon$1, whose $^1J_{\mathrm{CH}}$ deviates from the $^1J_{\mathrm{CH}}$/chemical shift correlation on which the adiabatic filter relies and is therefore not fully suppressed. In a 1:1 $^{13}$C:$^{12}$C mixture at the same total protein concentration (Supporting Figure S8A, right), the filtered spectrum retains ~40% of the unfiltered signal, consistent with selective retention of the $^1$H spins of the unlabeled component.

**Dependence of intermolecular NOEs on protein concentration.** To establish that the cross-peaks observed report on intermolecular contacts, two-dimensional $^1$H–$^1$H $^{13}$C-filtered/$^{13}$C-edited NOESY spectra were recorded on 1:1 $^{13}$C:$^{12}$C Tau-5* mixtures at total protein concentrations of 0.4, 0.6, 0.8 and 1.0 mM, in 100% $D_2O$ at 278 K (Supporting Figure S8B). Cross-peaks arising from incomplete filtering are intramolecular, so they would reproduce the same pattern at all concentrations, scaled only by the amount of labeled protein. In contrast, intermolecular NOEs depend on the population of contacts between labeled and unlabeled chains, which increases with protein concentration for a self-associating protein such as Tau-5*. Consistent with an intermolecular origin, the number and intensity of cross-peaks connecting $^{13}$C-bound aromatic protons (direct dimension, 6.2–7.5 ppm) with

$^{12}$C-bound protons of the unlabeled chains (indirect dimension), including methyl (∼0.8–1.2 ppm) and H$\alpha$ (∼4 ppm) protons, increase progressively from 0.4 to 1.0 mM. At 0.4 mM only a few weak cross-peaks are observed, whereas at 1.0 mM intense correlations to aliphatic and H$\alpha$ protons are detected. Together, these controls show that the $^{13}$C filter removes most of the $^{13}$C-bound magnetization and that the cross-peaks measured in solution samples report on intermolecular contacts between Tau-5* monomers.

#### Comparing NMR intermolecular NOE measurements with molecular dynamics simulations

To compare the simulations with experimental intermolecular NOEs, we used MDTraj[14] to calculate the average number of intermolecular proton contacts involving aromatic side-chain protons for which experimental NOEs were measured. For each trajectory frame, we measured pairwise distances between selected aromatic protons and all non-exchangeable protons from the opposite monomer. Non-exchangeable protons were considered to form an intermolecular contact with an aromatic side chain proton if they were within 6.0 Å.

Aromatic protons were grouped based on experimentally observed chemical shift overlap and analyzed in three merged categories:

- Trp H$\delta_1$: includes tryptophan aromatic protons at the H$\delta_1$ position.
- Tyr H$\delta_{1/2}$ or Trp H$\zeta_3$: includes tyrosine H$\delta_1$, H$\delta_2$ and tryptophan H$\zeta_3$ protons.
- Tyr H$\varepsilon_{1/2}$: includes tyrosine H$\varepsilon_1$ and H$\varepsilon_2$ protons.

For each class of aromatic side chain proton group, we computed the average number of intermolecular proton contacts observed over both monomers. We use the MD proton-contact counts as a qualitative structural metric rather than a quantitative back-calculation of NOE intensities, which would require $r^{-6}$ weighting and explicit treatment of dynamics.

We note that experimental NOE measurements were performed on the full 119-residue Tau-5* domain[36] and MD simulations were performed using a capped 24-residue Tau-5$_{\mathrm{R2}}$

fragment containing only the dominant oligomerization interface of Tau-5[31,37]. These values are therefore not expected to be directly quantitatively comparable. The increases in several intermolecular aromatic NOE signals are qualitatively consistent with the increased intermolecular aromatic proton contacts observed in the MD ensembles, although the experimental Tyr $\varepsilon$ signal changes little (**Figure 2D**).

#### Comparing NMR intermolecular chemical shift measurements with molecular dynamics simulations

Bielskutė-García et al.[31] determined protein-induced chemical shift perturbations (CSPs) by measuring chemical shifts in the Tau-5* region in the absence and presence of unlabeled $^{14}$N Tau-5*. These measurements were performed both in the absence and presence of EPI-001, yielding two sets of experimental protein-induced CSPs. To compare these experimental measurements with our MD simulation data, we used the SPARTA+[38] algorithm to compute $^{1}$H and $^{15}$N chemical shifts for each residue of both monomers in every frame of the apo and EPI-002 Tau-$5_{\mathrm{R2}}$ simulations. For each simulation condition, we calculated CSPs from the differences between average chemical shift values in dimer and monomer states. We compute MD-predicted CSPs using the same equation as Bielskutė-García et al.[31]:

$$\mathrm{CSP} = \left[ (\Delta\delta_{^{1}\mathrm{H}})^2 + \left( \frac{\Delta\delta_{^{15}\mathrm{N}}}{5} \right)^2 \right]^{1/2} . \tag{S3}$$

Here, $\Delta\delta_{^{1}\mathrm{H}}$ and $\Delta\delta_{^{15}\mathrm{N}}$ denote the differences between the average $^{1}$H and $^{15}$N chemical shifts in dimer and monomer states using the state definitions in "Molecular dynamics simulation analyses". Supporting Figure S3 compares the protein-induced CSPs obtained from MD simulations and experiments for the Tau-$5_{\mathrm{R2}}$ region under apo and ligand-containing conditions; the ligand-containing MD ensemble contains EPI-002, whereas the corresponding experimental measurements contain EPI-001. We note that EPI-001 contains a racemic mixture of four stereoisomers, previously shown to have similar affinities to Tau-5.[36] In this study we choose to simulate only EPI-002, the stereoisomer previously observed to have the highest affinity to Tau-5 by NMR spectroscopy.[36] Additionally, EPI-506, a clinical candidate for castration-resistant prostate cancer, is a prodrug of EPI-002, further motivating our choice to study this stereoisomer.

We observe reasonable agreement between the residue-wise patterns of CSPs calculated from the MD simulations and those measured experimentally upon dimerization and ternary complex formation, although the calculated CSPs are substantially larger in absolute magnitude. Quantitative agreement in CSP magnitude is not expected because chemical shift changes upon protein complex formation are often small relative to the intrinsic errors of structure-based chemical shift prediction methods[38,39]. We therefore interpret these comparisons primarily in terms of the locations and relative magnitudes of the residue-specific perturbations.

We compare the change in $\alpha$-helical propensity upon oligomerization in Supporting Figure S5. Experimental $\alpha$-helical propensities were obtained for monomeric Tau-5* and covalently linked Tau-5*[31] using $\delta$2D[40] propensities derived from $^{1}$H, $^{15}$N, C′, and C$\alpha$ chemical shifts measured from 2D $^{1}$H–$^{15}$N correlation spectra and 3D HNCO/HNCA experiments.[31] The experimental change in $\alpha$-helical propensity was computed by subtracting the monomeric Tau-5* dataset from the covalently linked Tau-5* dataset. For the MD simulations, each residue was assigned as $\alpha$-helical or non-$\alpha$-helical in every frame using the DSSP algorithm.[17] These assignments were averaged separately over the dimer and monomer frames to obtain the residue-specific ensemble-averaged $\alpha$-helical propensities. The MD-predicted change in Tau-$5_{R2}$ $\alpha$-helical content was then computed as the difference between the dimer and monomer ensemble-averaged $\alpha$-helical propensities. Positive values indicate increased $\alpha$-helical propensity in the dimeric or covalently linked state relative to the monomeric state.

Lastly, we compare residue-wise EPI-001-induced CSPs reported by Bielskutė-García et al.[31] with MD-estimated residue-wise EPI-002 contact probabilities in Supporting Figure S4. Experimental CSPs were determined by comparing NMR chemical shifts measured in the absence and presence of EPI-001 and calculated using Equation S3. For the MD simulations, the contact probability between each residue of Tau-$5_{R2}$ and EPI-002 was calculated as the fraction of frames in which the minimum distance between any heavy atom of EPI-002 and any heavy atom of the residue was less than 6 Å. Error bars for the MD-estimated contact probabilities were calculated by block averaging.

#### Consistency with previously reported AR mutational analyses

The molecular features associated with Tau-$5_{R2}$ oligomerization in our MD simulations are strongly supported by previously reported mutational studies of Tau-5* and the AR activation domain (Supporting Tables S2 and S3). The simulations identify a heterogeneous ensemble of intermolecular interfaces enriched in two recurring features: increased $\alpha$-helical propensity and intermolecular interactions involving aromatic side chains. Bielskuté-García et al. independently perturbed these properties in Tau-5*: A398P and L436P disrupt R2 and R3 helicity, respectively; G407A and G394A/S395A/G407A (AAA) increase R2 helical propensity; h2SA and h3A reduce aromatic character in R2 and R3, respectively; and C404Y increases R2 aromatic character with little effect on secondary structure.[31]

These perturbations altered both Tau-5* self-association and EPI-001 interaction propensity in the directions predicted by the simulations (Supporting Table S2). Concentration-dependent $^{1}H$–$^{15}N$ CSPs upon oligomerization increased when R2 helicity or aromatic character was increased and decreased when these features were disrupted. The same trend was observed for $^{1}H$–$^{15}N$ CSPs induced by one molar equivalent of EPI-001. For example, the strongly helix-stabilizing AAA mutant increased the mean R2 oligomerization CSP from 0.00995 to 0.02591 ppm and the mean R2 EPI-001-induced CSP from 0.00466 to 0.00986 ppm, whereas h2SA reduced these values to 0.00119 and 0.00154 ppm, respectively. EPI-001 condensate partitioning followed the same sequence dependence: relative to matched WT measurements, $P_{\text{condensate}}$ decreased by 47.2% for A398P and 69.1% for h2SA, but increased by 33.9% for G407A, 44.2% for AAA, and 99.8% for C404Y. Because the helicity and aromaticity partitioning series were measured under different solution conditions, all comparisons are made only to the matched WT condition within each series.[31]

These results connect the local R2/R3 interaction mechanism to condensate behavior. G407A, which increases R2 helicity without increasing aromatic content, and C404Y, which increases R2 aromaticity with comparatively little effect on secondary structure, both increased EPI-001 partitioning and produced less spherical, more arrested Tau-5* condensates. The rate of covalent modification of C404 by EPI-001 also correlated with Tau-5* helical and aromatic character and could not be explained by differences in intrinsic Cys nucleophilicity, consistent with prolonged ligand residence at the oligomerization-dependent interaction site.[31] Together, these measurements support the mechanism observed in our simulations, in which EPI-002 preferentially interacts with and stabilizes partially helical, aromatic-rich intermolecular Tau-$5_{\mathrm{R2}}$ interfaces.

Earlier experiments on the complete AR activation domain provide independent support for the same physical determinants (Supporting Table S3). Basu et al. progressively reduced aromatic valency using 8YtoS, 14YtoS, and 22YtoS variants. At 50 $\mu$M protein and 500 mM NaCl, WT* phase-separated at $T_C$ = 34 °C, but none of the aromatic mutants phase-separated below 60 °C. To observe phase separation of the weaker mutants, the authors increased both protein concentration to 150 $\mu$M and NaCl to 1 M; under these more favorable conditions, 8YtoS and 14YtoS had $T_C$ values of 31 and 48 °C, respectively, while 22YtoS still did not phase-separate. The mutant $T_C$ values therefore cannot be compared directly with the WT* value measured at lower protein and salt concentrations. EPI-001 partitioning decreased from approximately 55 in WT AR AD condensates to approximately 32 in 8YtoS condensates. Helix-breaking substitutions also showed a strong regional dependence: substitutions in Tau-5 increased $T_C$ by approximately 10 °C and reduced condensate number and size in cells, whereas analogous substitutions in Tau-1 increased $T_C$ by only approximately 5 °C and had little cellular effect. Conversely, increasing AR AD helical propensity with 2.5% and 5% TFE decreased $T_C$ by 12 and 35 °C, respectively.[28]

Together, the two experimental studies show that transient helical structure and aromatic interactions promote Tau-5/AR self-association and condensation and that the same molecular features promote EPI-001 interaction and condensate enrichment. These independent perturbations therefore support the ensemble-level mechanism observed in our simulations, in which ligand binding stabilizes dynamic Tau-$5_{\mathrm{R2}}$ oligomerization interfaces through multivalent interactions involving partially helical, aromatic-rich regions.

**Supporting Table S2. Tau-5* mutational analysis reported by Bielskuté-García et al.**[31] Concentration-dependent oligomerization CSPs are mean $^{1}$H–$^{15}$N CSPs induced by increasing Tau-5* concentration from 25 to 400 $\mu$M. EPI-001-induced CSPs are mean $^{1}$H–$^{15}$N CSPs measured for 250 $\mu$M Tau-5* upon addition of 250 $\mu$M EPI-001 (one molar equivalent). CSPs were calculated from the $^{1}$H and $^{15}$N chemical-shift differences using Equation S3; regional means were then calculated over R2 (residues 391–414) and R3 (residues 426–446), excluding residues without evaluable CSPs. $P_{\text{condensate}}$ denotes the EPI-001 dense/light-phase partition coefficient. Partitioning was measured with 3 mg/mL Tau-5* at 500 mM NaCl and 250 $\mu$M EPI-001 for the helicity series, and at 1.75 M NaCl and 100 $\mu$M EPI-001 for the aromaticity series.

| **Tau-5* variant** | **Perturbation** | **Oligomerization-induced Tau-5* $^{1}$H–$^{15}$N CSPs R2 / R3 (ppm)** | **EPI-001-induced Tau-5* $^{1}$H–$^{15}$N CSPs R2 / R3 (ppm)** | **EPI-001 $P_{\text{condensate}}$** | **Experimental result** |
|---|---|---|---|---|---|
| **Helicity series** | | | | | |
| WT (3 mg/mL Tau-5*; 500 mM NaCl; 250 $\mu$M EPI-001) | Native reference | 0.0100 / 0.0055 | 0.0047 / 0.0036 | $22.56 \pm 1.99$ | Reference. |
| A398P | Disrupts R2 helix | 0.0046 / 0.0036 | 0.0026 / 0.0022 | $11.91 \pm 0.45$ (−47.2% vs WT) | Reduced R2 self-association and EPI-001 enrichment. |
| L436P | Disrupts R3 helix | 0.0070 / 0.0033 | 0.0031 / 0.0019 | $12.91 \pm 0.78$ (−42.8%) | Reduced R3 self-association and EPI-001 enrichment. |
| G407A | Stabilizes R2 helix | 0.0159 / 0.0091 | 0.0062 / 0.0044 | $30.21 \pm 1.27$ (+33.9%) | Increased self-association, EPI-001 enrichment, and aspect ratio. |
| AAA | Stabilizes R2 helix (G394A/S395A/G407A) | 0.0259 / 0.0109 | 0.0099 / 0.0062 | $32.53 \pm 1.56$ (+44.2%) | Increased self-association and EPI-001 enrichment. |
| **Aromaticity series** | | | | | |
| WT (3 mg/mL Tau-5*; 1.75 M NaCl; 100 $\mu$M EPI-001) | Native reference | 0.0100 / 0.0055 | 0.0047 / 0.0036 | $161.11 \pm 9.04$ | Reference. |
| h2SA | Reduces R2 aromaticity (Y393S/W397A/Y406S/H413A) | 0.0012 / 0.0023 | 0.0015 / 0.0007 | $49.81 \pm 1.45$ (−69.1% vs WT) | Reduced R2 self-association and EPI-001 enrichment. |
| h3A | Reduces R3 aromaticity (W433A/H434A/F437A) | 0.0062 / 0.0027 | 0.0018 / 0.0007 | $71.47 \pm 16.27$ (−55.6%) | Reduced R3 self-association and EPI-001 enrichment. |
| C404Y | Increases R2 aromaticity; little secondary-structure change | 0.0219 / 0.0091 | 0.0062 / 0.0042 | $321.87 \pm 12.73$ (+99.8%) | Increased self-association and EPI-001 enrichment without increased helicity; increased aspect ratio. |
| noAro | Removes aromatic residues | 0.0017 / 0.0012 | 0.0012 / 0.0007 | — | Reduced self-association and EPI-001 interaction. |

**Supporting Table S3. AR activation-domain mutational analysis reported by Basu et al.**[28] The table summarizes independent perturbations of aromatic character and transient helicity and their effects on AR phase separation and EPI-001 partitioning. The aromatic-mutant cloud-point experiments used 50 $\mu$M protein and 500 mM NaCl for WT* ($T_C$ = 34 °C); none of the YtoS mutants phase-separated below 60 °C under these conditions. To resolve the mutant phase transitions, both protein concentration and salt were raised to 150 $\mu$M protein and 1 M NaCl: 8YtoS and 14YtoS then had $T_C$ = 31 °C and 48 °C, respectively, whereas 22YtoS still did not phase-separate. These $T_C$ values should not be compared directly across the two conditions. $P_{\mathrm{EPI-001}}$ denotes the EPI-001 dense/light-phase partition coefficient.

| **Construct / perturbation** | **Intended structural change** | **Experimental result** |
|---|---|---|
| AR AD WT* | Reference construct (L26P background for in vitro phase-separation experiments) | $T_C$ and EPI-001 partitioning were measured; $P_{\mathrm{EPI-001}} \approx 55$. |
| 8YtoS | Reduced aromatic character | $T_C = 31$ °C under mutant-resolving conditions; $P_{\mathrm{EPI-001}} \approx 32$, approximately 40% below WT. |
| 14YtoS | Further reduced aromatic character | $T_C = 48$ °C under mutant-resolving conditions. |
| 22YtoS | Removes all Tyr residues | No phase separation under mutant-resolving conditions; reduced partitioning into MED1-IDR/RNAPII-CTD condensates and no DHT-induced condensates. |
| L56P | Helix-breaking substitution in the polyQ region | $T_C$ increased by approximately 5 °C. |
| Tau-1 A186P/L192P/C238P | Disrupts transient Tau-1 helices | $T_C$ increased by approximately 5 °C; little effect on cellular condensation. |
| Tau-5 A356P/A398P/T435P | Disrupts transient Tau-5 helices, including the R2/R3 region | $T_C$ increased by approximately 10 °C; condensate number and size decreased in cells. |
| WT* + 2.5% TFE | Increases AR AD helical propensity | $T_C$ decreased by 12 °C. |
| WT* + 5% TFE | Further increases AR AD helical propensity | $T_C$ decreased by 35 °C. |

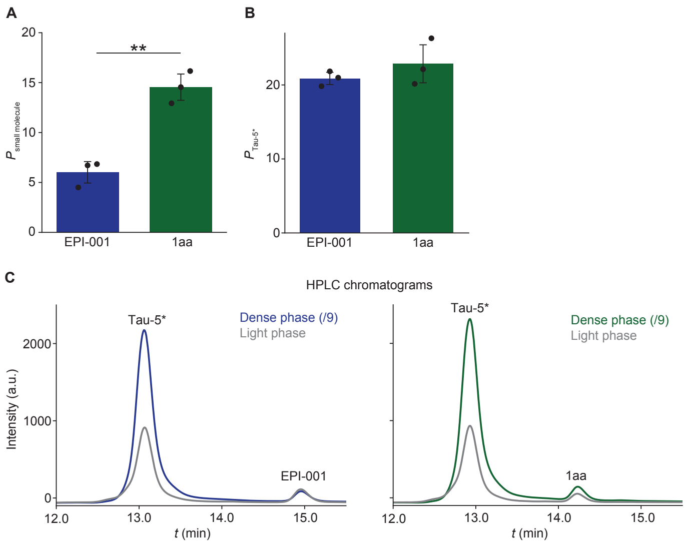


**Supporting Figure S1. Partitioning of EPI-001 and 1aa into Tau-5* condensates.** **(A)** EPI-001 and 1aa partition coefficients into Tau-5* condensates. **(B)** Tau-5* partition coefficient in the presence of EPI-001 and 1aa. **(C)** HPLC chromatograms showing Tau-5*, EPI-001 and 1aa signals in the dense and light phases of the protein. Dense-phase samples were diluted ninefold prior to HPLC analysis. Measurements were performed in triplicate. Bars represent the mean of three measurements, and error bars indicate standard deviations calculated using divisor $n$. Double asterisks (**) indicate $P < 0.01$.

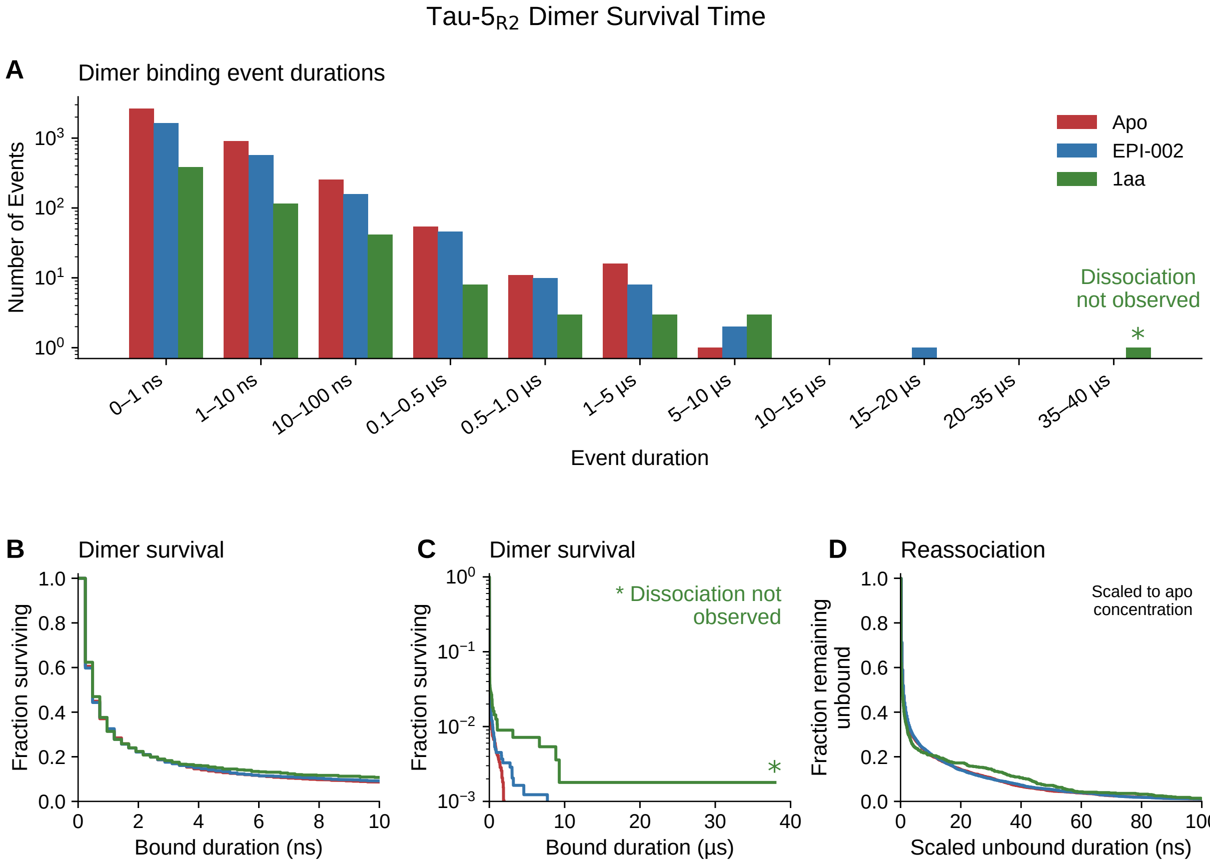


**Supporting Figure S2. Dimer survival and concentration-scaled reassociation of Tau-5$_{R2}$.** (**A**) Distribution of observed dimer binding-event durations in the absence of ligands (red) and in the presence of EPI-002 (blue) or 1aa (green). (**B–C**) Fraction of binding events remaining associated as a function of time since association, shown over the initial 10 ns on a linear scale (B) and over the full duration range with a logarithmic vertical axis (C). Asterisks indicate the terminal 1aa binding event, for which dissociation was not observed. (**D**) Fraction of events remaining unbound as a function of time since dissociation, scaled to the mean apo partner concentration (3.327 mM) and shown over 0–100 ns on a linear time axis. Frames were classified as associated when at least one intermolecular C$\alpha$–C$\alpha$ minimum-image distance was below 10 Å. Event durations were resolved at 0.24-ns intervals. Survival estimates account for events ongoing at the trajectory endpoint; concentration scaling and event definitions are described in "Dimer survival and concentration-scaled reassociation."

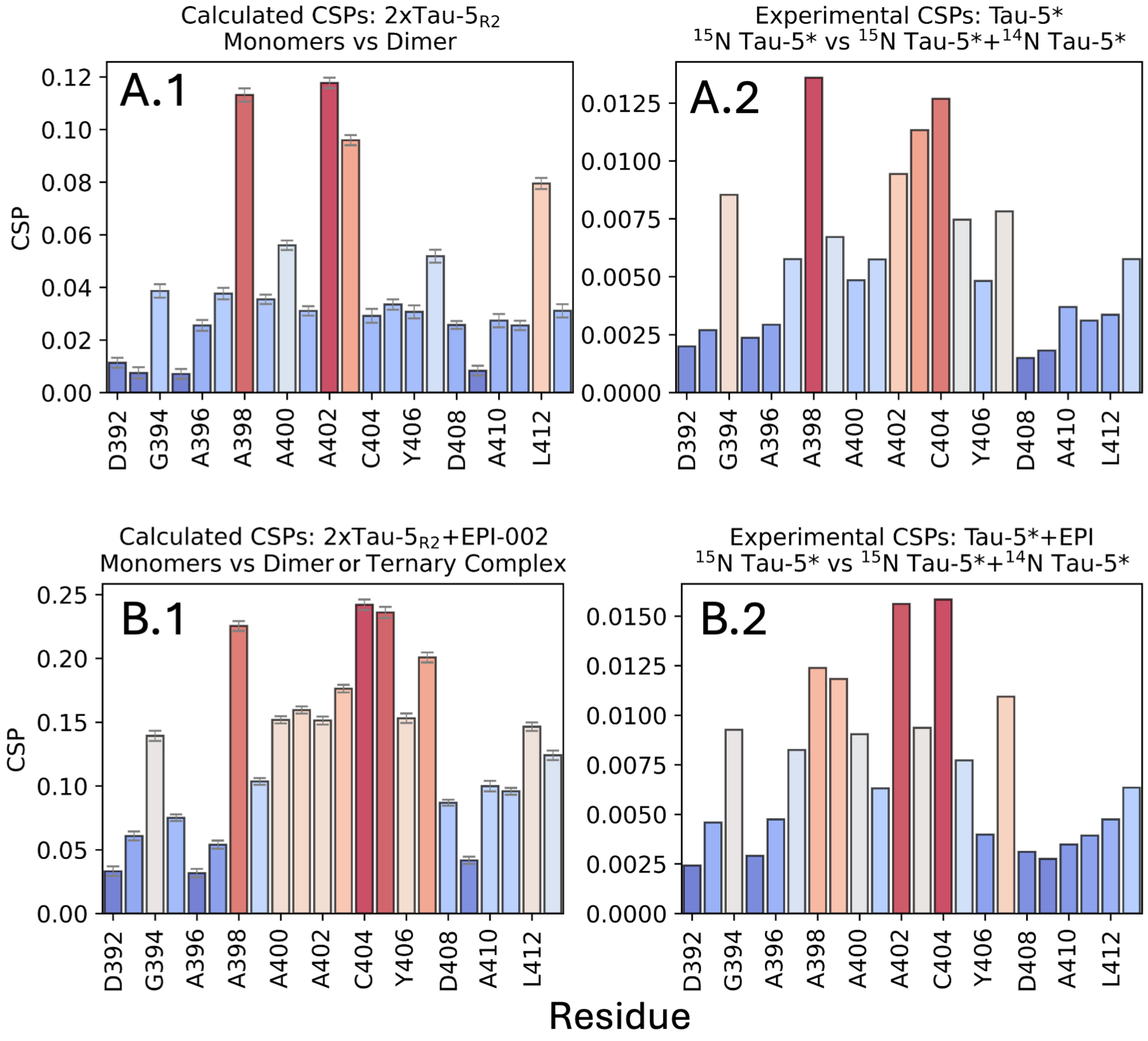


**Supporting Figure S3. Comparison of MD-predicted and experimental protein-induced $^{1}$H–$^{15}$N chemical shift perturbations (CSPs) under apo and ligand-containing conditions.** (**A.1**) MD-predicted Tau-5$_{R2}$ $^{1}$H–$^{15}$N CSPs obtained by computing the perturbation between dimer and monomer ensemble-averaged chemical shifts. (**A.2**) $^{1}$H–$^{15}$N CSPs over the same Tau-5$_{R2}$ residues reported by Bielskutė-García *et al.*[31], obtained by measuring chemical shifts of the Tau-5* region in the presence and absence of unlabeled $^{14}$N Tau-5*. (**B.1**) The corresponding MD-predicted protein-induced $^{1}$H–$^{15}$N CSPs in the presence of EPI-002. (**B.2**) Experimental protein-induced $^{1}$H–$^{15}$N CSPs measured in the presence of EPI-001. Bars are colored by CSP magnitude within each panel, from blue (smaller perturbations) through white to red (larger perturbations).

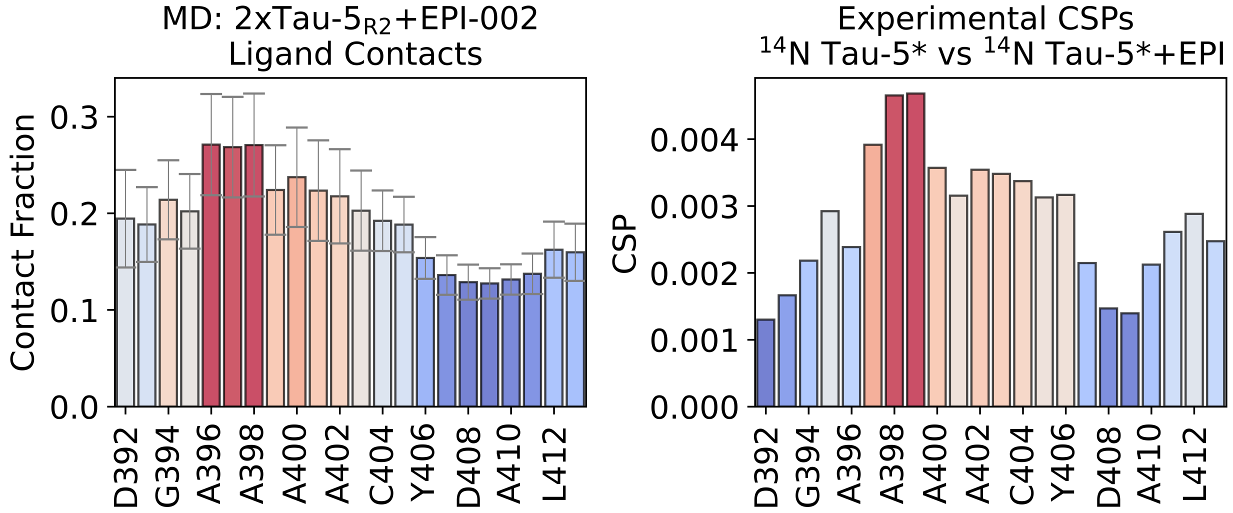


**Supporting Figure S4. Comparison of MD-predicted protein–EPI-002 interactions and experimental ligand-induced $^{1}$H–$^{15}$N chemical shift perturbations (CSPs).** (**A**) Contact probabilities between each residue of the R2 domain and EPI-002 observed in an unbiased 100 $\mu$s MD simulation performed with the a99SB-disp force field. A contact was assigned to all MD frames in which the minimum distance between any heavy atom of EPI-002 and any heavy atom of a protein residue was $< 6$ Å. Error bars were calculated by block averaging. (**B**) Experimental $^{1}$H–$^{15}$N CSPs of the R2 domain of Tau-5* induced by EPI-001 binding, calculated by comparing NMR chemical shifts measured in the absence and presence of EPI-001[31]. Bars are colored by the magnitude of the plotted quantity within each panel, from blue (lower values) through white to red (higher values): contact probability in panel A and CSP magnitude in panel B.

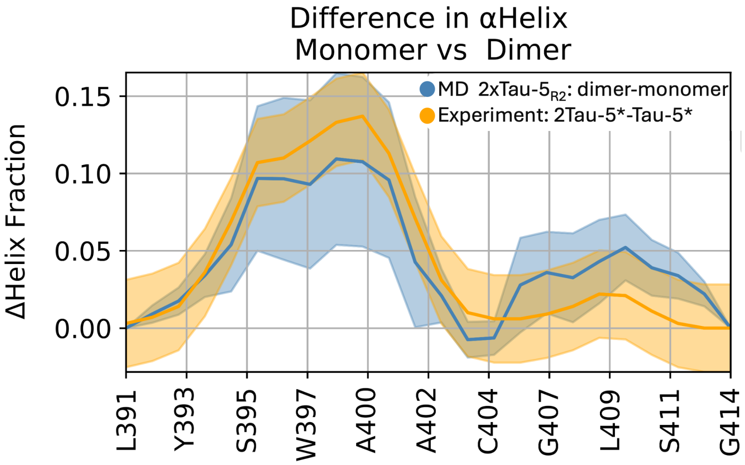


**Supporting Figure S5. Comparison of MD-predicted and experimentally measured changes in Tau-5 $\alpha$-helical propensity upon oligomerization.** (blue) MD-predicted Tau-$5_{R2}$ change in $\alpha$-helical propensity, computed as the difference between dimer and monomer ensemble-averaged $\alpha$-helical propensities assigned using the DSSP algorithm. (orange) Experimental change in Tau-5* $\alpha$-helical propensity, computed as the $\alpha$-helical propensity of covalently linked Tau-5* minus that of monomeric Tau-5* using $\delta$2D[40] propensities derived from $^{1}$H, $^{15}$N, C′, and C$\alpha$ chemical shifts measured from 2D $^{1}$H–$^{15}$N correlation spectra and 3D HNCO/HNCA experiments. Positive values indicate increased $\alpha$-helical propensity in the dimeric or covalently linked state relative to the monomeric state. The agreement between the MD-predicted and experimentally inferred changes supports the conclusion that Tau-5 oligomerization promotes local $\alpha$-helical structure within the Tau-$5_{R2}$ region.

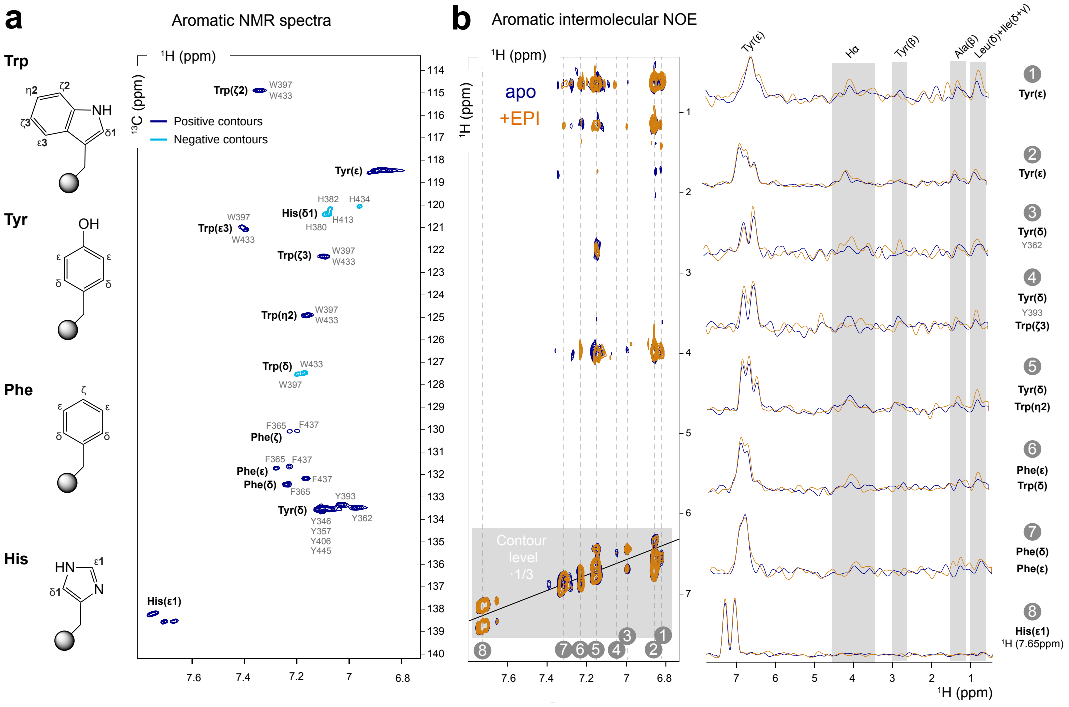


**Supporting Figure S6. Intermolecular NOE measurements for apo and EPI-001–bound Tau-5*.** (a) Two-dimensional $^{1}$H–$^{13}$C TROSY spectrum of the aromatic region. Positive contours are shown in blue and negative contours in cyan; these colors indicate contour sign, not different sample conditions. (b) Overlaid two-dimensional $^{1}$H–$^{1}$H NOE spectra and corresponding one-dimensional traces showing intermolecular contacts between isotopically labeled and unlabeled monomers in the absence (blue) and presence (orange) of EPI-001.

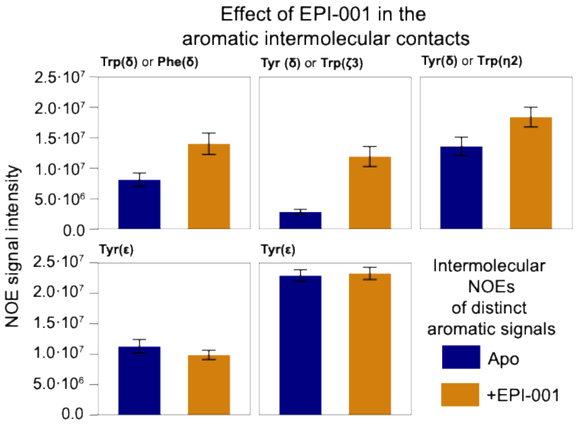


**Supporting Figure S7. Experimental intermolecular NOE intensities for Tau-5* dimers.** Intermolecular NOE intensities measured from samples containing a mixture of $^{13}$C,$^{1}$H-labeled and $^{12}$C,$^{1}$H-labeled Tau-5* in the absence (blue) and presence (orange) of EPI-001, recorded at 800 MHz using the $^{13}$C-filtered/$^{13}$C-edited pulse scheme of Zwahlen *et al.*[33]. Peaks correspond to aromatic protons as assigned from NMR spectra.

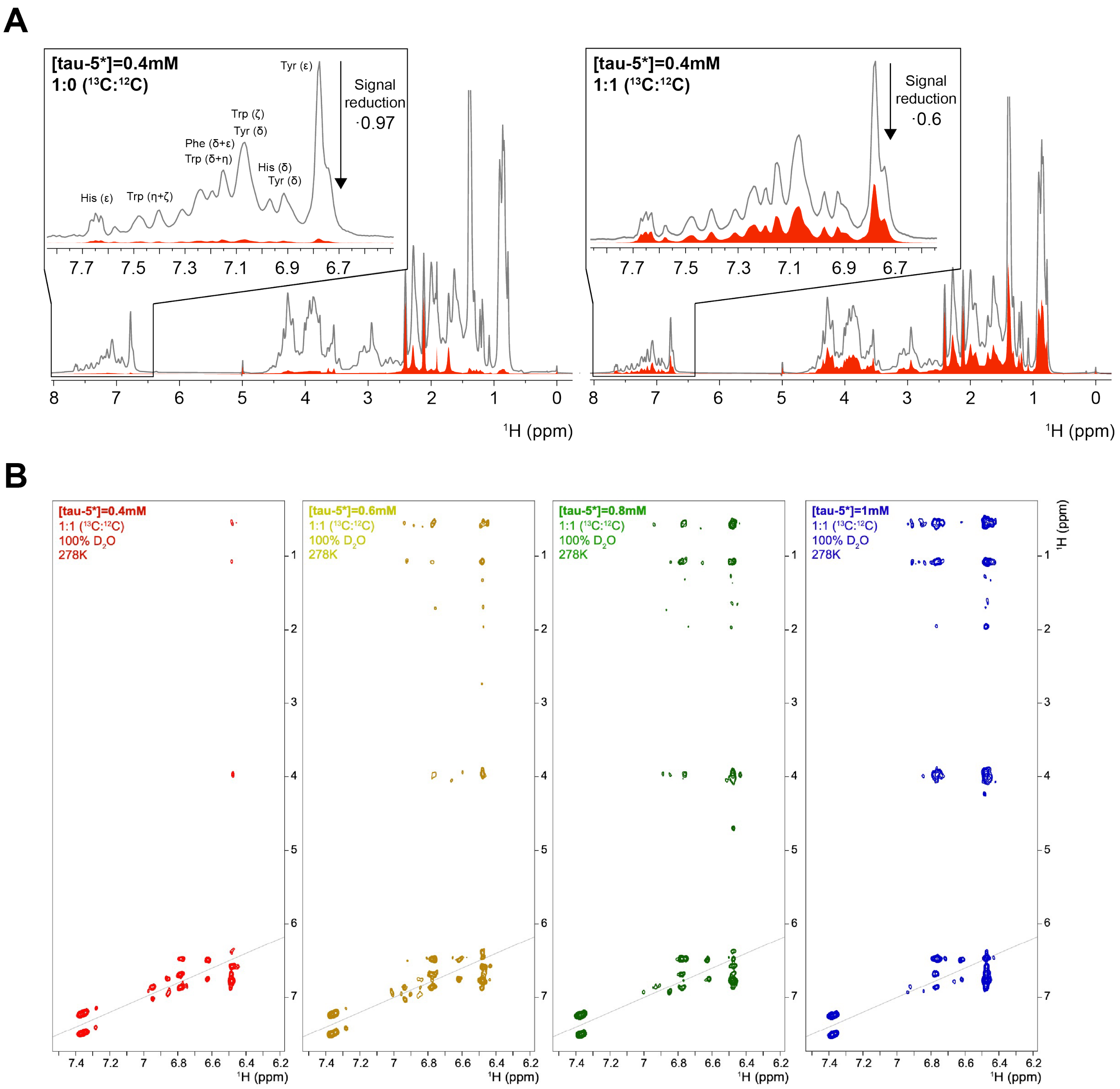


**Supporting Figure S8. Controls for the measurement of intermolecular NOEs in Tau-5*.** (A) Efficiency of the $^{13}$C filter. One-dimensional $^1$H spectra of 0.4 mM Tau-5* recorded without (grey) and with (red) the double $^{13}$C filter of the $^{13}$C-filtered/$^{13}$C-edited NOESY pulse scheme of Zwahlen *et al.*,[33] for uniformly $^{13}$C-labeled Tau-5* (1:0 $^{13}$C:$^{12}$C, left) and for a 1:1 mixture of $^{13}$C,$^1$H- and $^{12}$C,$^1$H-labeled Tau-5* (right). Insets show expansions of the aromatic region with resonance assignments. The filter suppresses ∼97% of the signal in the uniformly $^{13}$C-labeled sample. In the 1:1 mixture, ∼40% of the unfiltered signal is retained, corresponding to the unlabeled component. (B) Two-dimensional $^1$H–$^1$H $^{13}$C-filtered/$^{13}$C-edited NOESY spectra of 1:1 $^{13}$C:$^{12}$C Tau-5* mixtures at total protein concentrations of 0.4 (red), 0.6 (yellow), 0.8 (green) and 1.0 mM (blue), recorded at 800 MHz in 100% $D_2O$ at 278 K. The number and intensity of intermolecular cross-peaks between $^{13}$C-bound aromatic protons and $^{12}$C-bound protons increase with protein concentration.

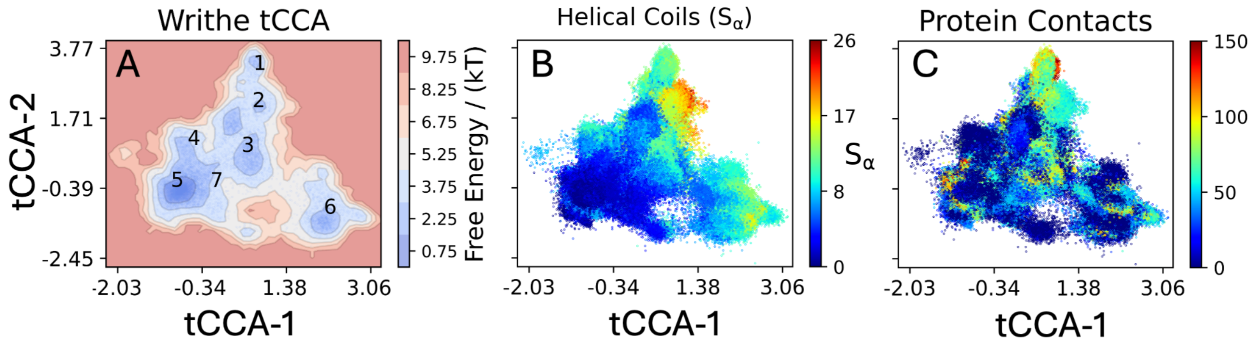


**Supporting Figure S9. Time-lagged canonical correlation analysis (tCCA) of Tau-5$_{\mathrm{R2}}$ self-association in the absence of ligands using writhe structural descriptors.** (**A**) The free energy surface of Tau-5$_{\mathrm{R2}}$ conformational states as a function of the two slowest dynamic modes obtained from tCCA of writhe structural descriptors, with the centroid of each MSM state marked with its index. Additional molecular features are projected onto the same tCCA coordinates, with colors indicating (**B**) helical content quantified by the $S_\alpha$ order parameter and (**C**) the total number of intermolecular C$\alpha$–C$\alpha$ contacts.

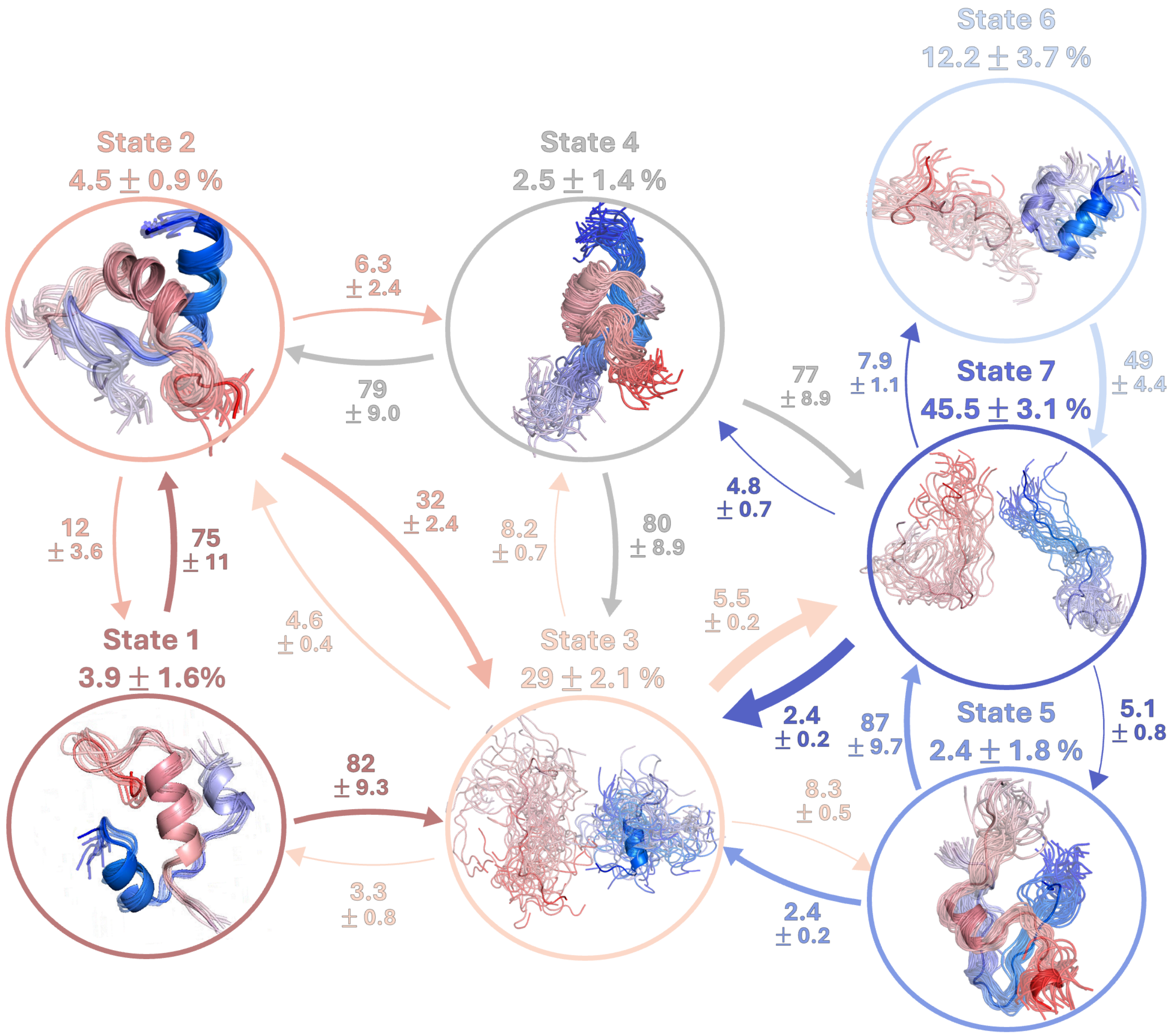


**Supporting Figure S10. Markov state model (MSM) of Tau-5$_{\mathrm{R2}}$ self-association in the absence of ligands estimated from an unbiased 100 $\mu$s molecular dynamics simulation.** Representative structures of each Markov state are displayed in circles with their steady-state probabilities above. The Tau-5$_{\mathrm{R2}}$ monomers are colored with a blue-to-red gradient from the N-terminus of the first monomer to the C-terminus of the second. Directed arrows indicate transition probability fluxes between states, with arrow thickness proportional to flux magnitude. Arrows are labeled with mean first passage times between states, reported in $\mu$s. Reported uncertainties are the average of the upper and lower deviations from the estimate to the bounds of the 95% confidence interval, calculated from 1,000 bootstrap samples.

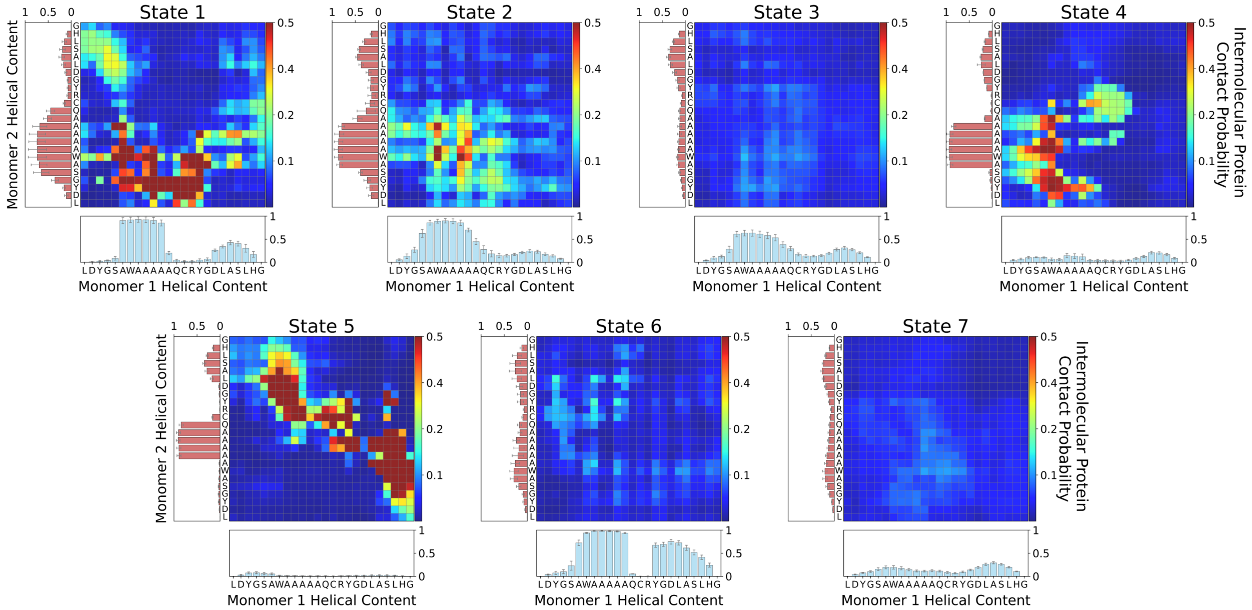


**Supporting Figure S11.** Intermolecular C$\alpha$–C$\alpha$ contact probabilities and residue-specific $\alpha$-helical populations (bottom and left) in each metastable conformational state of the MSM of Tau-$5_{\mathrm{R2}}$ self-association in the absence of ligands.

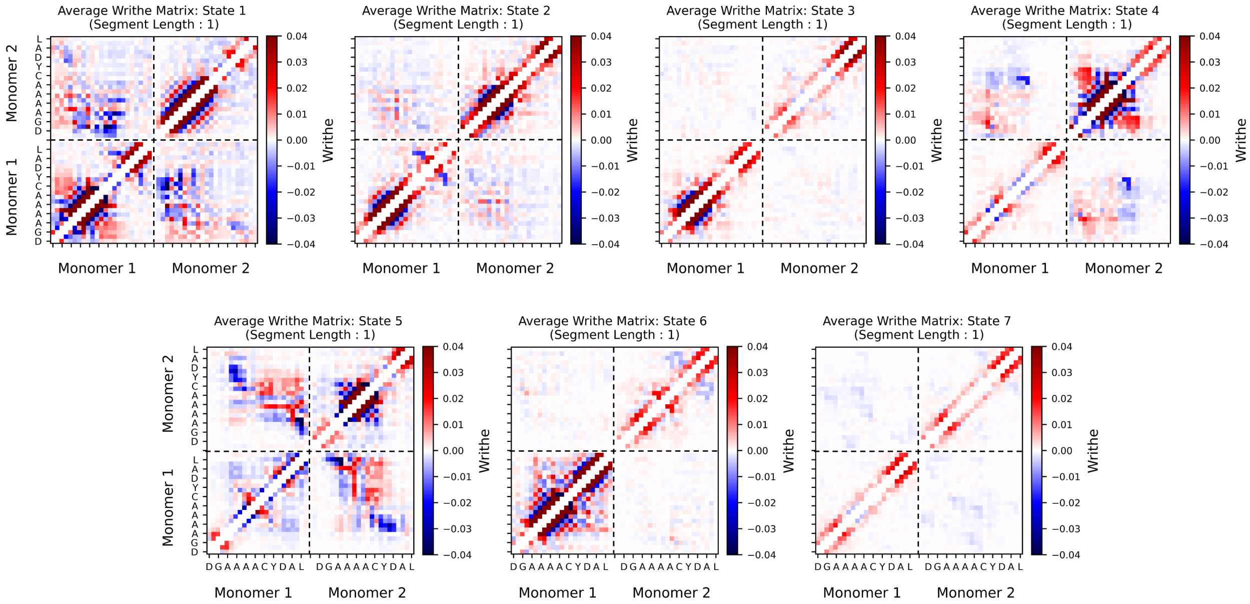


**Supporting Figure S12.** Average intramolecular (bottom left and top right quadrants) and intermolecular (top left and bottom right quadrants) writhe matrices in each metastable conformational state of the MSM of Tau-$5_{\mathrm{R2}}$ self-association in the absence of ligands. Each matrix entry shows the average signed crossing contribution of two protein backbone segments, each defined by a vector from the C$\alpha$ atom of residue $i$ to the C$\alpha$ atom of residue $i+1$.

**Supporting Table S4.** Average conformational properties of Tau-$5_{R2}$ states in the absence of ligands (apo). Each state corresponds to a kinetically distinct conformational state obtained from coarse-graining the apo Tau-$5_{R2}$ MSM using the generalized PCCA spectral clustering algorithm[26,27]. Reported quantities include the steady-state population obtained from the MSM (*Population %*), average number of intermolecular protein–protein contacts (*Protein Contacts*), helical content order parameter ($S\alpha$), and percentage of conformations forming dimers (*Dimer %*).

| State | Population % | Protein Contacts | $S\alpha$ | Dimer % |
|---|---|---|---|---|
| 1 | 3.9 | 101.1 | 12.8 | 99.8 |
| 2 | 4.5 | 66.3 | 14.2 | 98.3 |
| 3 | 28.8 | 37.8 | 7.9 | 70.3 |
| 4 | 2.5 | 52.8 | 5.5 | 84.1 |
| 5 | 2.4 | 89.3 | 3.6 | 99.5 |
| 6 | 12.2 | 31.5 | 10.4 | 61.8 |
| 7 | 45.5 | 31.2 | 3.0 | 56.9 |

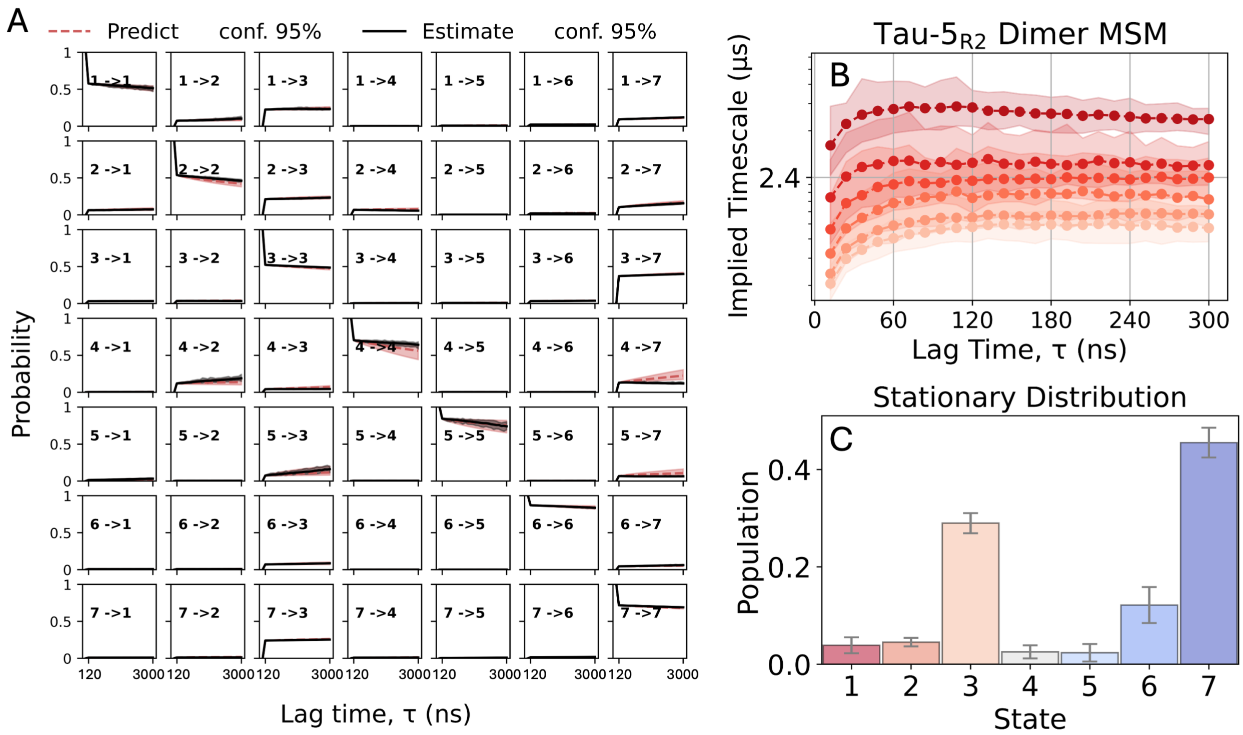


**Supporting Figure S13. Validation tests and stationary distribution for the MSM of Tau-5$_{R2}$ self-association in the absence of ligands.** (A) Chapman–Kolmogorov (CK) test comparing predicted and directly estimated transition probabilities between states $i$ and $j$ at integer multiples of the model lag time (120 ns). Red dotted lines ("Predict") show transition probabilities predicted by propagating the transition matrix while solid black lines ("Estimate") show transition probabilities obtained from transition matrices estimated directly from the trajectory data at integer multiples of the lag time. Red and black shaded regions indicate the 95% confidence interval of the mean obtained from Gibbs sampling with 1000 samples. (B) Log-scaled implied timescales (ITS) obtained from MSM transition matrices estimated at increasing lag times. Converged implied timescales are expected to be approximately independent of lag time. Colored shaded regions indicate the 95% confidence intervals of the mean implied timescales at each lag time. Here, all ITS are well above the threshold that can be resolved by the model. (C) Stationary probabilities of the MSM states, with error bars indicating the 95% confidence interval of the mean.

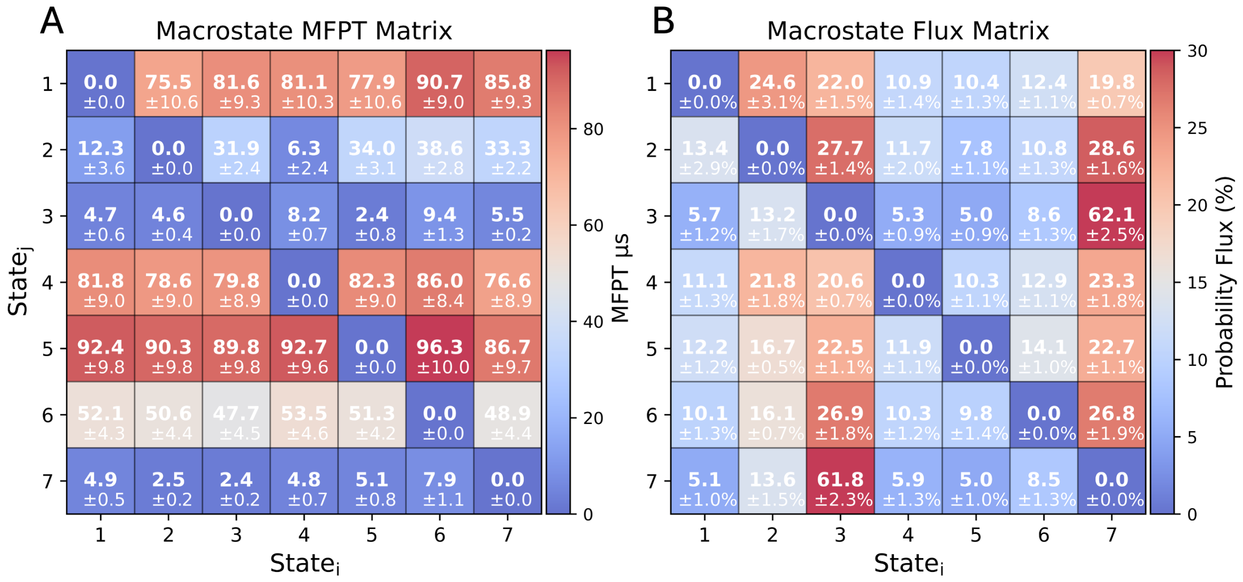


**Supporting Figure S14. Mean first-passage times and transition probability fluxes for the MSM of Tau-5$_{\mathrm{R2}}$ self-association in the absence of ligands.** (A) Mean first-passage times (MFPTs) and (B) row-normalized transition probability fluxes between metastable macrostates obtained by generalized PCCA spectral clustering of an MSM containing 150 microstates.

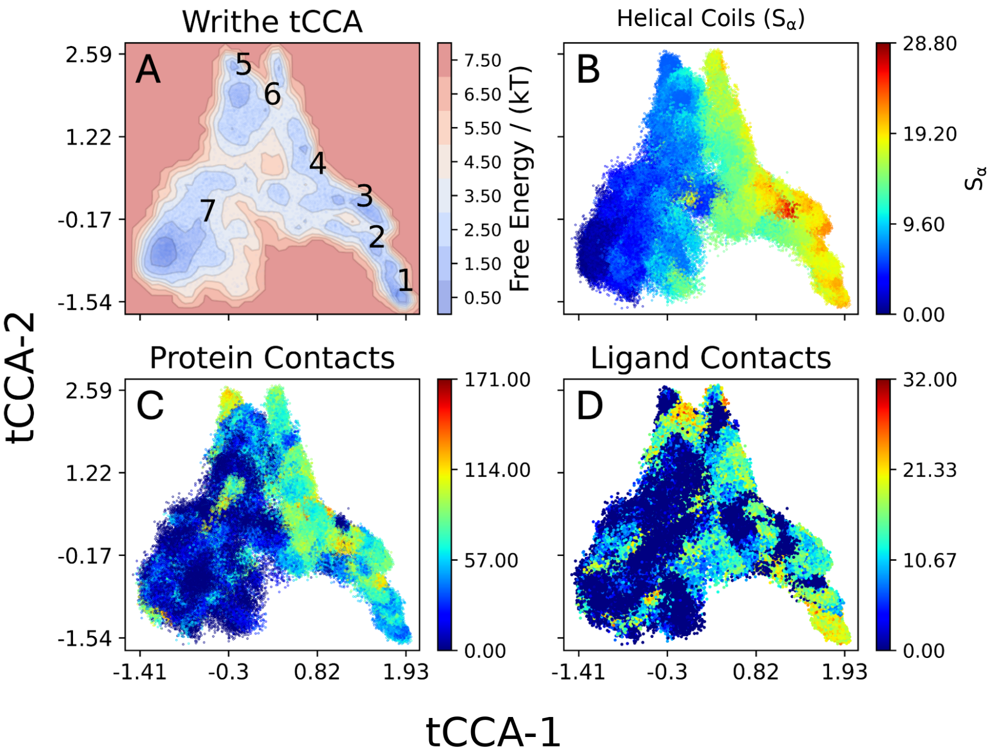


**Supporting Figure S15. Time-lagged canonical correlation analysis (tCCA) of Tau-5$_{R2}$ self-association in the presence of EPI-002 using writhe structural descriptors.** (**A**) The free energy surface of Tau-5$_{R2}$ conformational states as a function of the two slowest dynamic modes obtained from tCCA of intramolecular and intermolecular writhe features, with the centroid of each MSM state marked with its index. Additional molecular features are projected onto the same tCCA coordinates, with colors indicating (**B**) helical content quantified by the $S_\alpha$ order parameter, (**C**) the total number of intermolecular C$\alpha$–C$\alpha$ contacts, and (**D**) the total number of protein–ligand contacts.

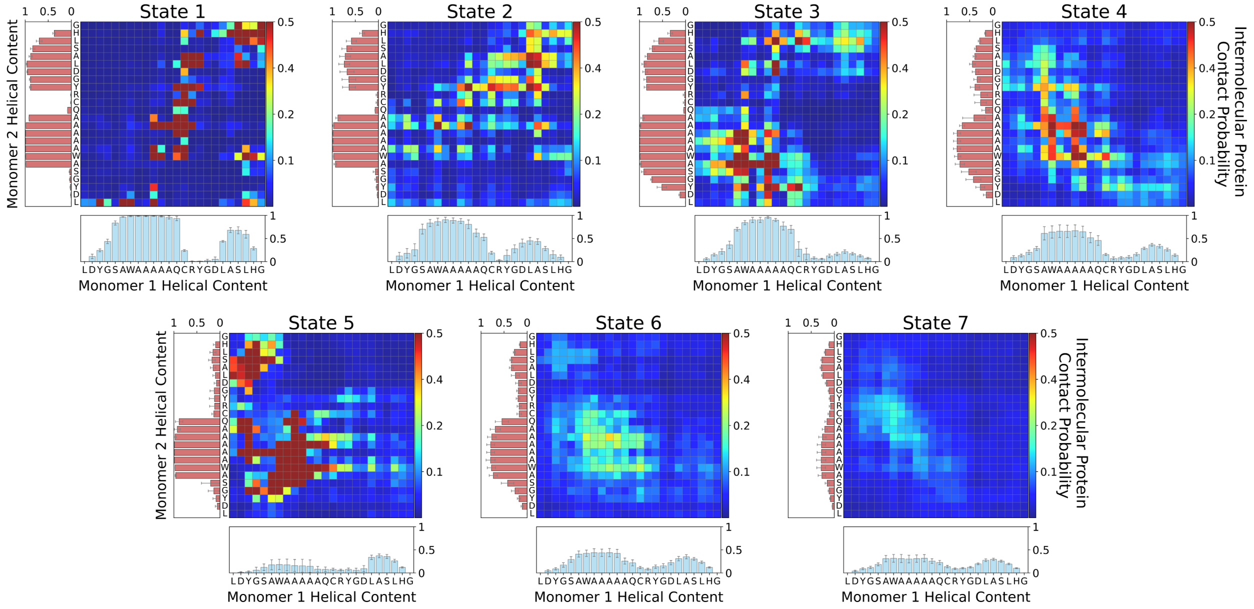


**Supporting Figure S16.** Intermolecular C$\alpha$–C$\alpha$ contact probabilities and residue-specific $\alpha$-helical populations (bottom and left) in each metastable conformational state of the MSM of Tau-$5_{\mathrm{R2}}$ self-association in the presence of EPI-002.

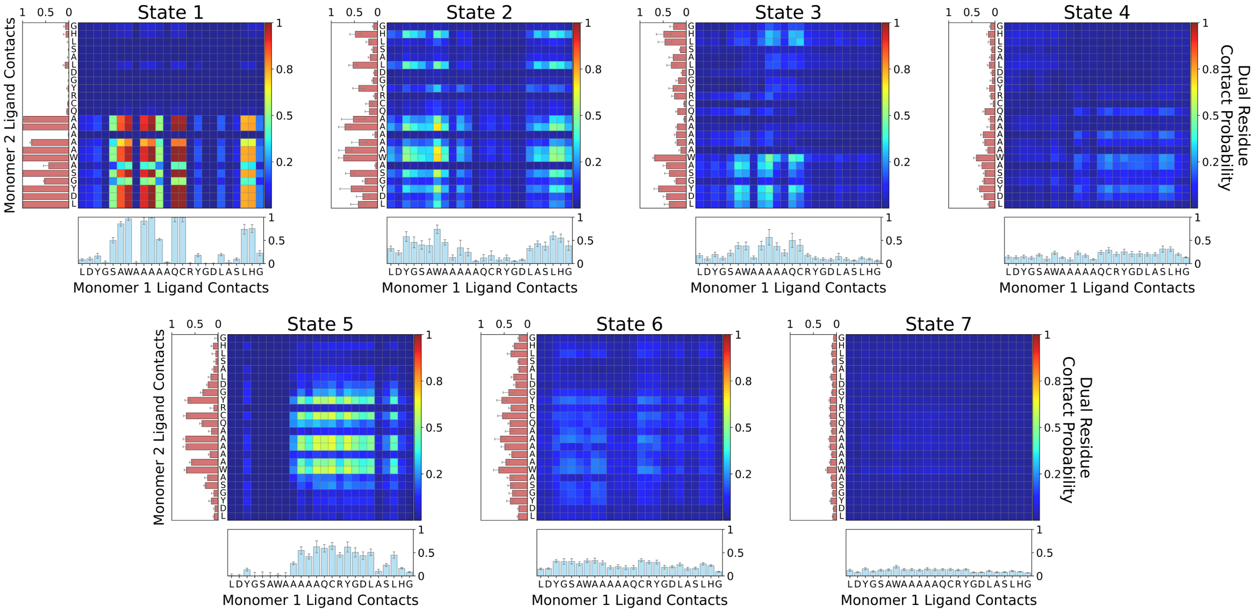


**Supporting Figure S17.** Probability that two residues, one from each Tau-$5_{\mathrm{R2}}$ monomer, simultaneously contact EPI-002 in each metastable conformational state of the MSM of Tau-$5_{\mathrm{R2}}$ self-association in the presence of EPI-002. Bar plots below (blue) and to the left (red) show the probabilities that individual residues in monomers 1 and 2, respectively, contact EPI-002.

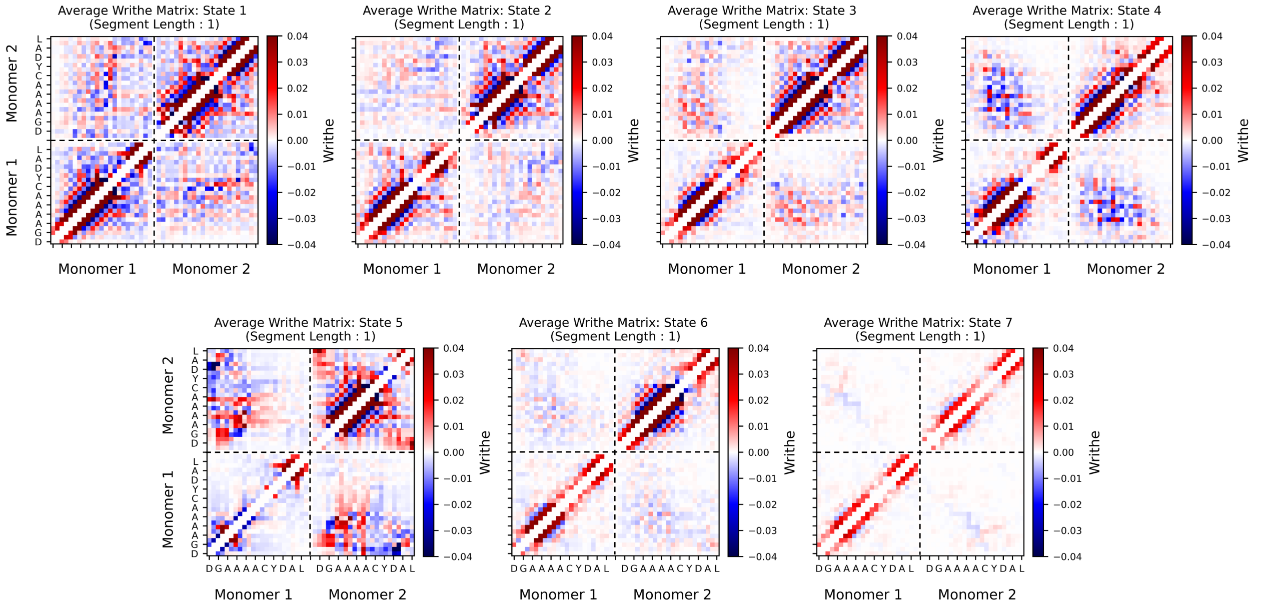


**Supporting Figure S18.** Average intramolecular (bottom left and top right quadrants) and intermolecular (top left and bottom right quadrants) writhe matrices in each metastable conformational state of the MSM of Tau-$5_{R2}$ self-association in the presence of EPI-002. Each matrix entry shows the average signed crossing contribution of two protein backbone segments, each defined by a vector from the C$\alpha$ atom of residue $i$ to the C$\alpha$ atom of residue $i + 1$.

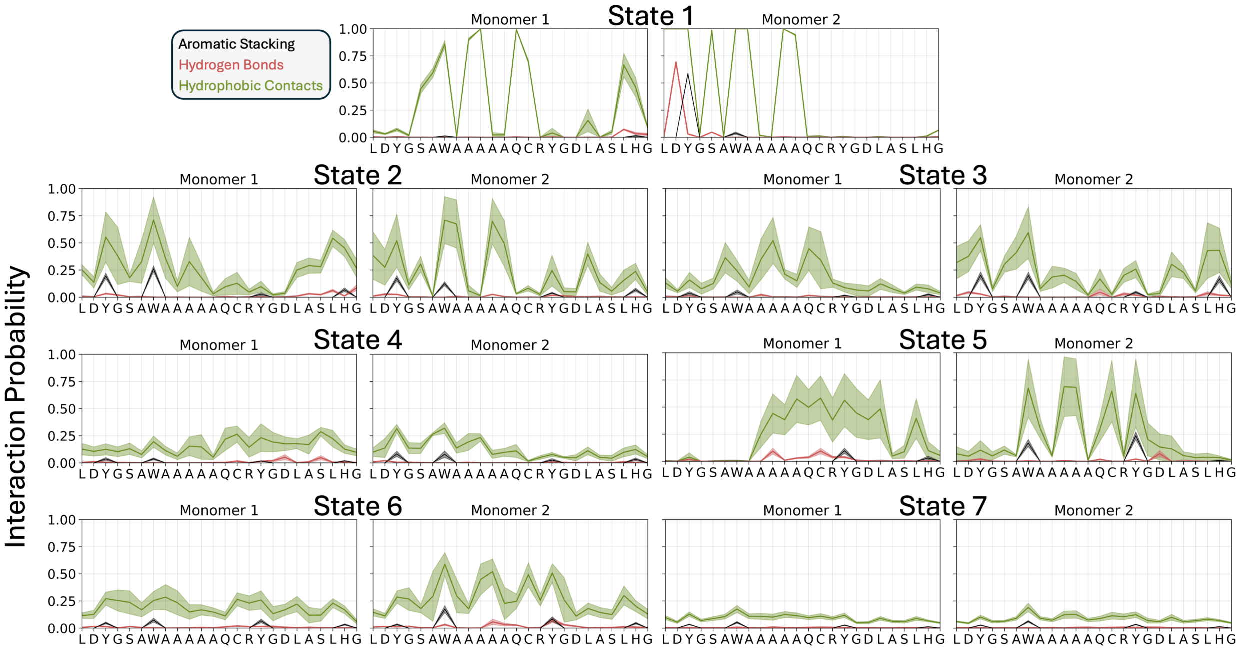


**Supporting Figure S19.** Probabilities of aromatic stacking (black), hydrogen bonding (red), and hydrophobic contacts (green) between EPI-002 and individual residues of both Tau-$5_{R2}$ monomers in each metastable conformational state of the MSM of Tau-$5_{R2}$ self-association in the presence of EPI-002.

**Supporting Table S5.** Average conformational properties of Tau-$5_{R2}$ metastable macrostates in the presence of EPI-002. Metastable states were obtained from an initial, fine-grained K-means clustering of writhe-based topological features computed from the EPI-002 + Tau-$5_{R2}$ simulation ensemble projected onto the 3 slowest dynamical modes identified by tCCA followed by MSM estimation and coarse-graining by generalized PCCA spectral clustering. Reported quantities include cluster population (*Population %*), average number of intermolecular protein–protein contacts (*Protein Contacts*), helical content order parameter ($S\alpha$), percentage of conformations forming dimers (*Dimer %*), average number of protein–ligand contacts (*Ligand Contacts*), and percentage of conformations forming ternary complexes (*Ternary %*).

| State | Population % | Protein Contacts | $S\alpha$ | Dimer % | Ligand Contacts | Ternary % |
|---|---|---|---|---|---|---|
| 1 | 7.4 | 53.0 | 18.3 | 99.5 | 18.8 | 97.4 |
| 2 | 8.0 | 53.4 | 15.9 | 95.8 | 14.7 | 80.4 |
| 3 | 8.0 | 66.8 | 17.8 | 96.2 | 12.2 | 78.0 |
| 4 | 8.7 | 60.7 | 11.5 | 83.9 | 9.0 | 51.6 |
| 5 | 3.4 | 81.1 | 8.4 | 96.5 | 13.6 | 70.6 |
| 6 | 16.2 | 49.0 | 9.8 | 81.9 | 11.1 | 54.6 |
| 7 | 48.3 | 35.3 | 5.1 | 64.2 | 5.3 | 26.6 |

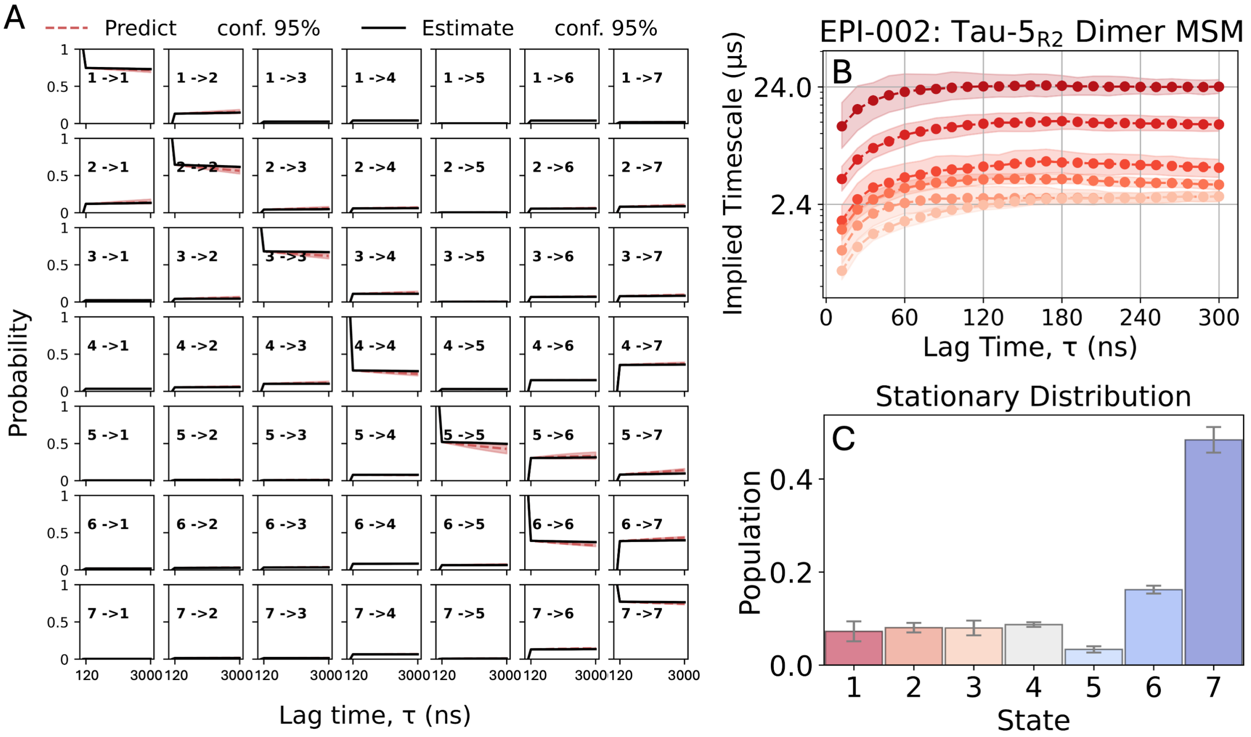


**Supporting Figure S20. Validation tests and stationary distribution for the MSM of Tau-5$_{R2}$ self-association in the presence of EPI-002.** (A) Chapman–Kolmogorov (CK) test comparing predicted and directly estimated transition probabilities between states $i$ and $j$ at integer multiples of the model lag time (120 ns). Red dotted lines ("Predict") show transition probabilities predicted by propagating the transition matrix while solid black lines ("Estimate") show transition probabilities obtained from transition matrices estimated directly from the trajectory data at integer multiples of the lag time. Red and black shaded regions indicate the 95% confidence interval of the mean obtained from Gibbs sampling with 1000 samples. (B) Log-scaled implied timescales (ITS) obtained from MSM transition matrices estimated at increasing lag times. Converged implied timescales are expected to be approximately independent of lag time. Colored shaded regions indicate the 95% confidence intervals of the mean implied timescales at each lag time. Here, all ITS are well above the threshold that can be resolved by the model. (C) Stationary probabilities of the MSM states, with error bars indicating the 95% confidence interval of the mean.

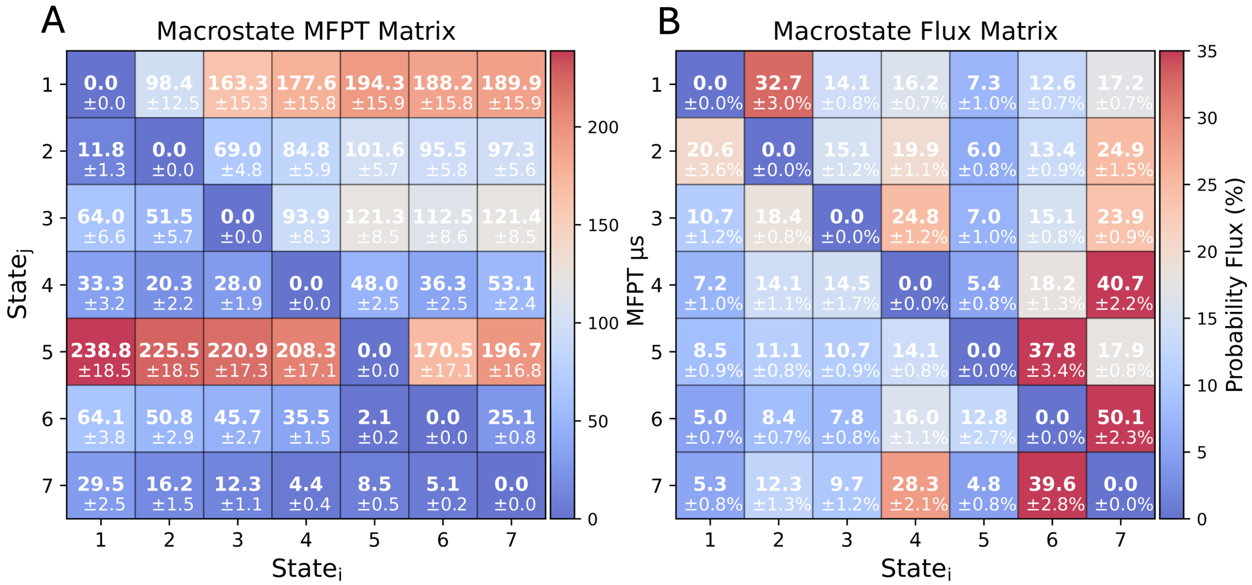


**Supporting Figure S21. Mean first-passage times and transition probability fluxes for the MSM of Tau-5$_{\mathrm{R2}}$ self-association in the presence of EPI-002.** (A) Mean first-passage times (MFPTs) and (B) row-normalized transition probability fluxes between metastable macrostates obtained by generalized PCCA spectral clustering of an MSM containing 400 microstates.

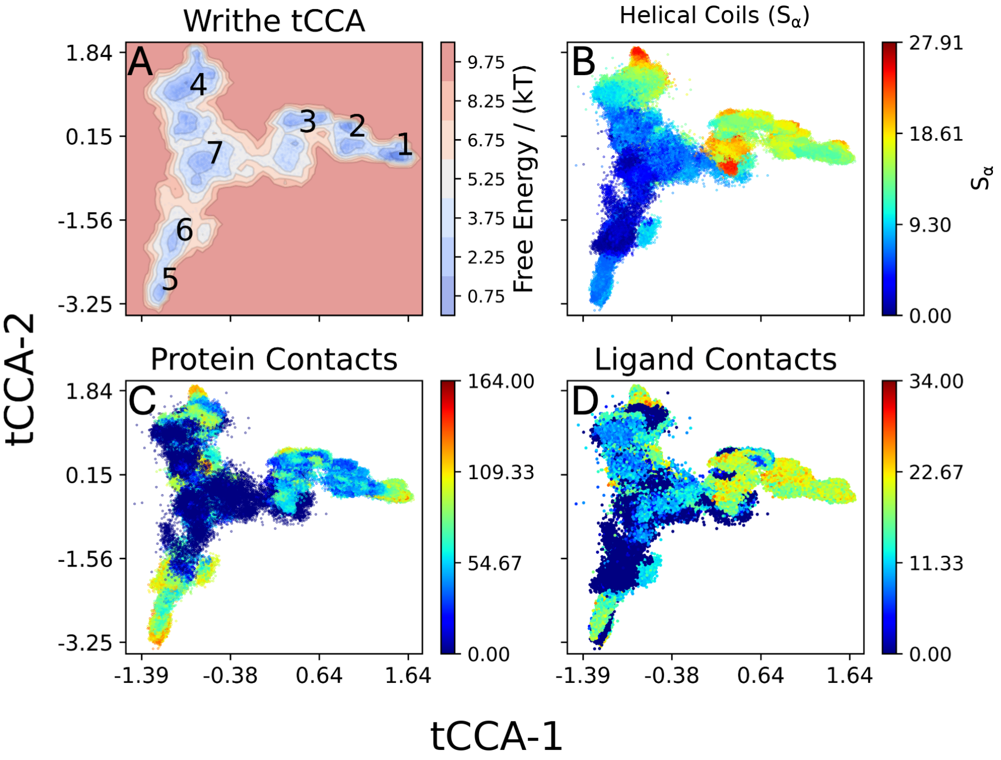


**Supporting Figure S22. Time-lagged canonical correlation analysis (tCCA) of Tau-5$_{R2}$ self-association in the presence of 1aa using writhe structural descriptors.** (**A**) The free energy surface of Tau-5$_{R2}$ conformational states as a function of the two slowest dynamic modes obtained from tCCA of intramolecular and intermolecular writhe features, with the centroid of each MSM state marked with its index. Additional molecular features are projected onto the same tCCA coordinates, with colors indicating (**B**) helical content quantified by the $S_\alpha$ order parameter, (**C**) the total number of intermolecular C$\alpha$–C$\alpha$ contacts, and (**D**) the total number of protein–ligand contacts.

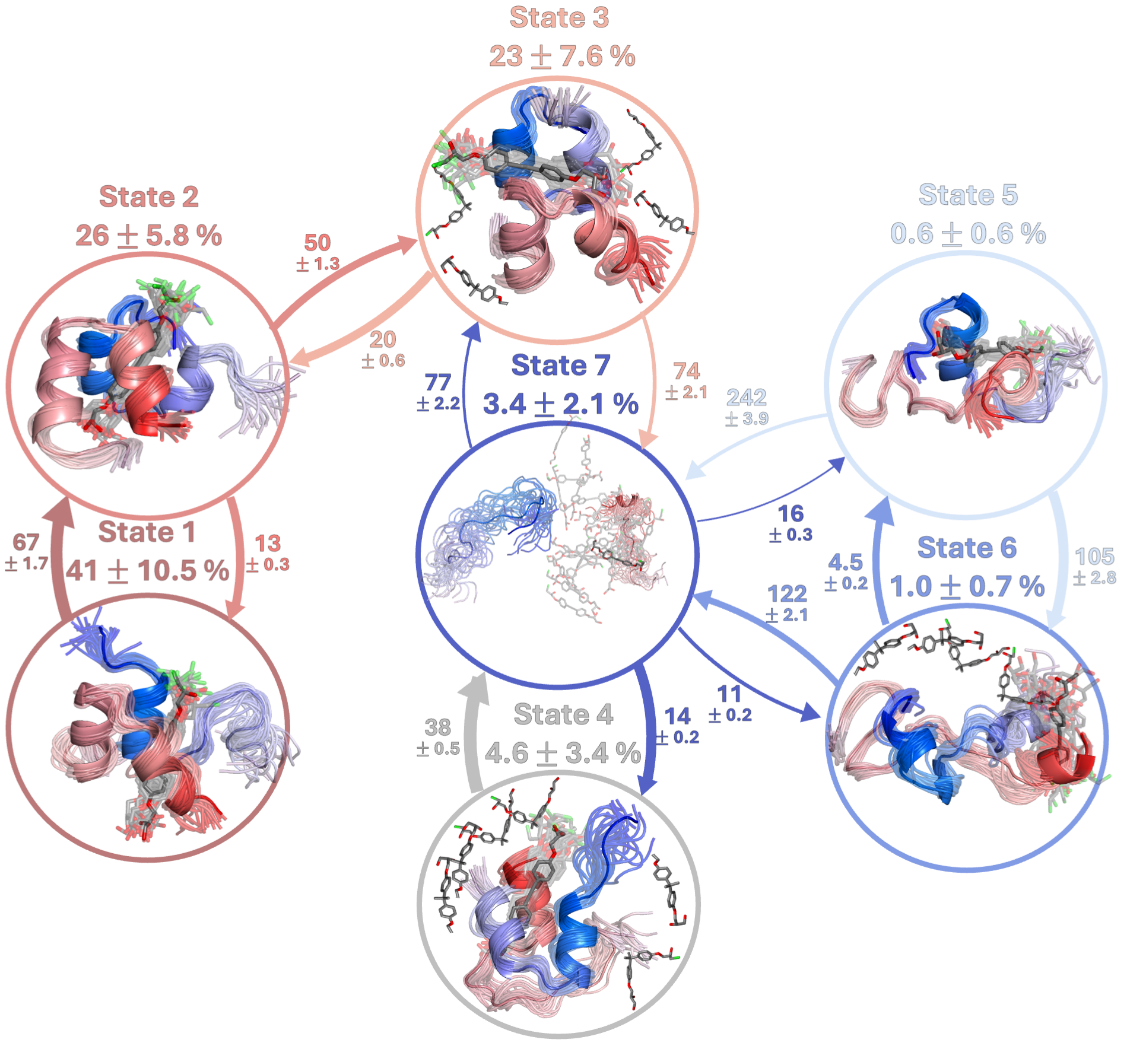


**Supporting Figure S23. Markov state model (MSM) of Tau-$5_{R2}$ self-association in the presence of 1aa estimated from an unbiased 90 $\mu$s molecular dynamics simulation.** Representative structures of each conformational state are displayed in circles with their steady-state probabilities above. The Tau-$5_{R2}$ monomers are colored with a blue-to-red gradient from the N-terminus of the first monomer to the C-terminus of the second. Directed arrows indicate transition probability fluxes between states, with arrow thickness proportional to flux magnitude. Arrows are labeled with mean first passage times between states, reported in $\mu$s. Reported uncertainties indicate the 95% confidence interval of the mean, calculated from 1,000 bootstrap samples.

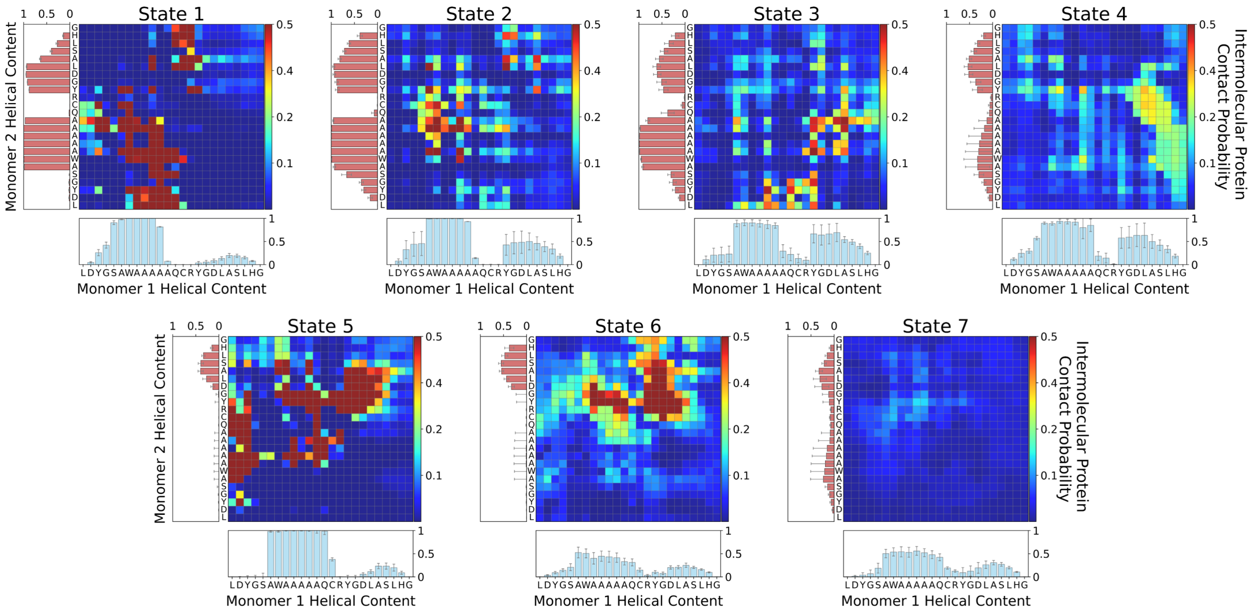


**Supporting Figure S24.** Intermolecular Cα–Cα contact probabilities and residue-specific α-helical populations (bottom and left) in each metastable conformational state of the MSM of Tau-$5_{R2}$ self-association in the presence of 1aa.

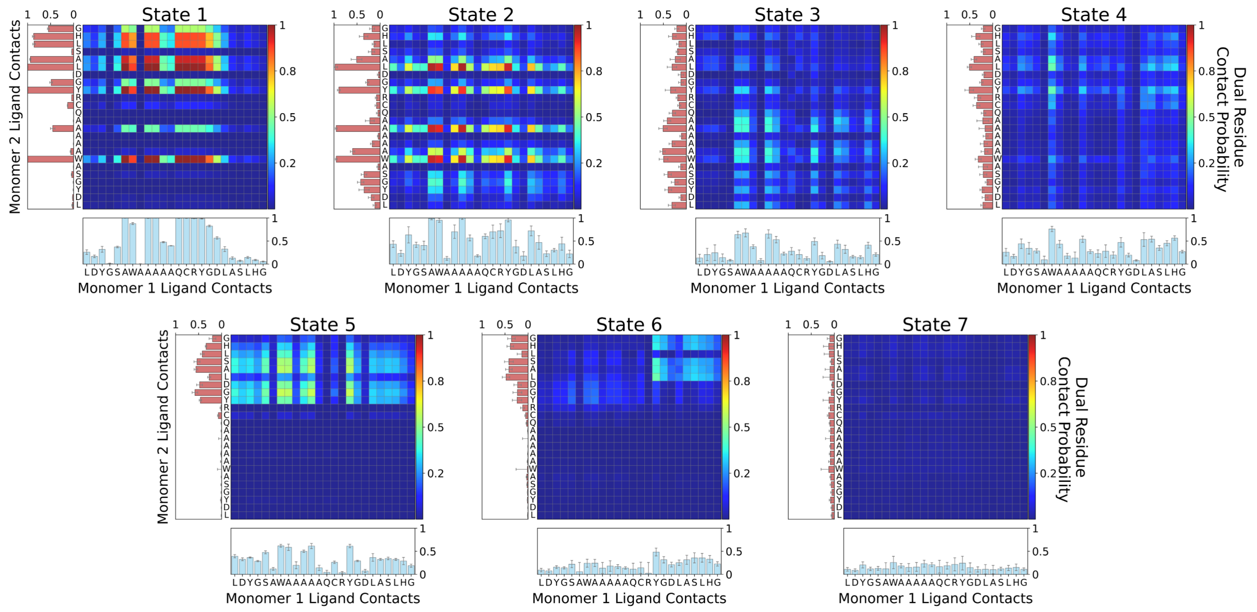


**Supporting Figure S25.** Probability that two residues, one from each Tau-$5_{R2}$ monomer, simultaneously contact 1aa in each metastable conformational state of the MSM of Tau-$5_{R2}$ self-association in the presence of 1aa. Bar plots below (blue) and to the left (red) show the probabilities that individual residues in monomers 1 and 2, respectively, contact 1aa.

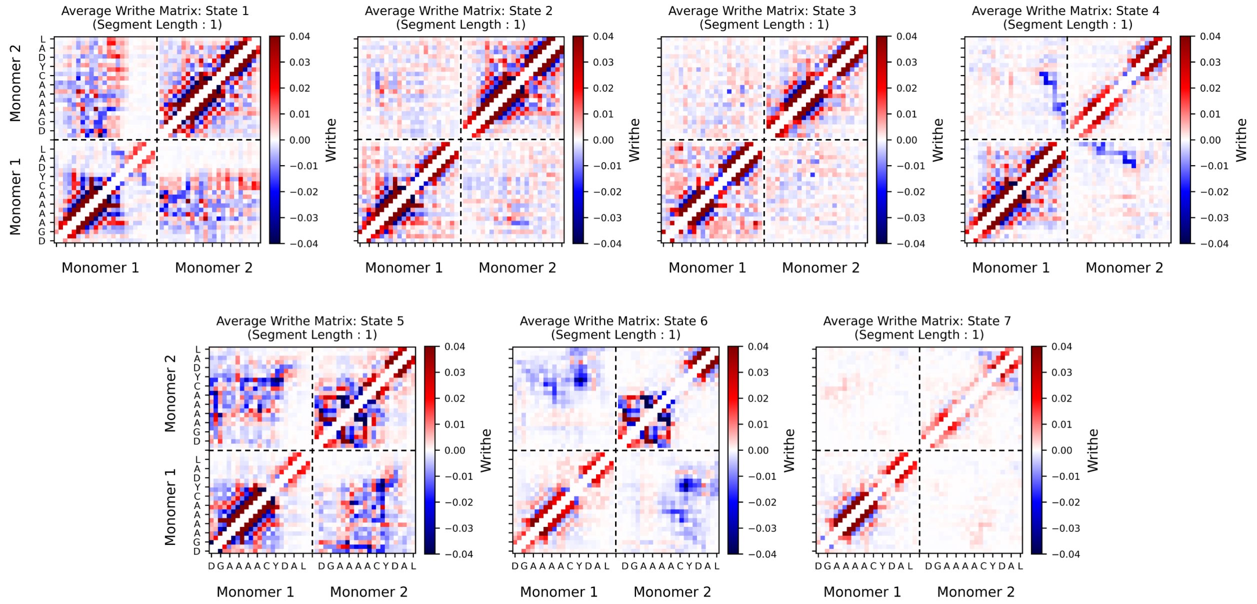


**Supporting Figure S26.** Average intramolecular (bottom left and top right quadrants) and intermolecular (top left and bottom right quadrants) writhe matrices in each metastable conformational state of the MSM of Tau-$5_{R2}$ self-association in the presence of 1aa. Each matrix entry shows the average signed crossing contribution of two protein backbone segments, each defined by a vector from the C$\alpha$ atom of residue $i$ to the C$\alpha$ atom of residue $i + 1$.

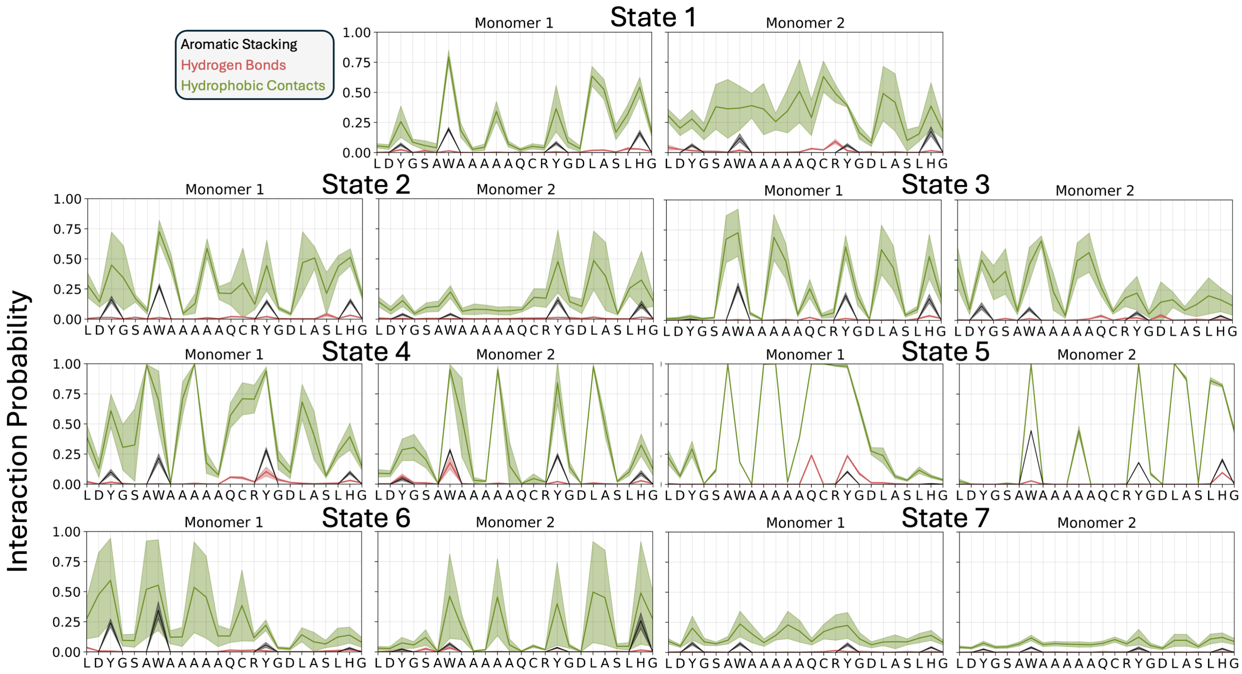


**Supporting Figure S27.** Probabilities of aromatic stacking (black), hydrogen bonding (red), and hydrophobic contacts (green) between 1aa and individual residues of both Tau-$5_{R2}$ monomers in each metastable conformational state of the MSM of Tau-$5_{R2}$ self-association in the presence of 1aa.

**Supporting Table S6.** Average conformational properties of Tau-$5_{R2}$ metastable macrostates in the presence of 1aa. Metastable states were obtained from an initial, fine-grained K-means clustering of writhe-based topological features computed from the 1aa + Tau-$5_{R2}$ simulation ensemble projected onto the 3 slowest dynamical modes identified by tCCA followed by MSM estimation and coarse-graining by generalized PCCA spectral clustering. Reported quantities include cluster population (*Population %*), average number of intermolecular protein–protein contacts (*Protein Contacts*), helical content order parameter ($S\alpha$), percentage of conformations forming dimers (*Dimer %*), average number of protein–ligand contacts (*Ligand Contacts*), and percentage of conformations forming ternary complexes (*Ternary %*).

| State | Population % | Protein Contacts | $S\alpha$ | Dimer % | Ligand Contacts | Ternary % |
|---|---|---|---|---|---|---|
| 1 | 41.2 | 83.1 | 15.0 | 99.9 | 19.6 | 99.8 |
| 2 | 25.7 | 50.8 | 17.4 | 99.5 | 19.6 | 97.9 |
| 3 | 23.0 | 52.2 | 16.0 | 88.5 | 14.4 | 78.7 |
| 4 | 4.9 | 55.1 | 12.8 | 80.7 | 14.8 | 74.0 |
| 5 | 0.6 | 102.9 | 7.9 | 98.8 | 12.0 | 63.9 |
| 6 | 1.0 | 76.3 | 6.7 | 96.1 | 10.1 | 64.5 |
| 7 | 3.6 | 40.5 | 10.1 | 64.6 | 11.3 | 49.9 |

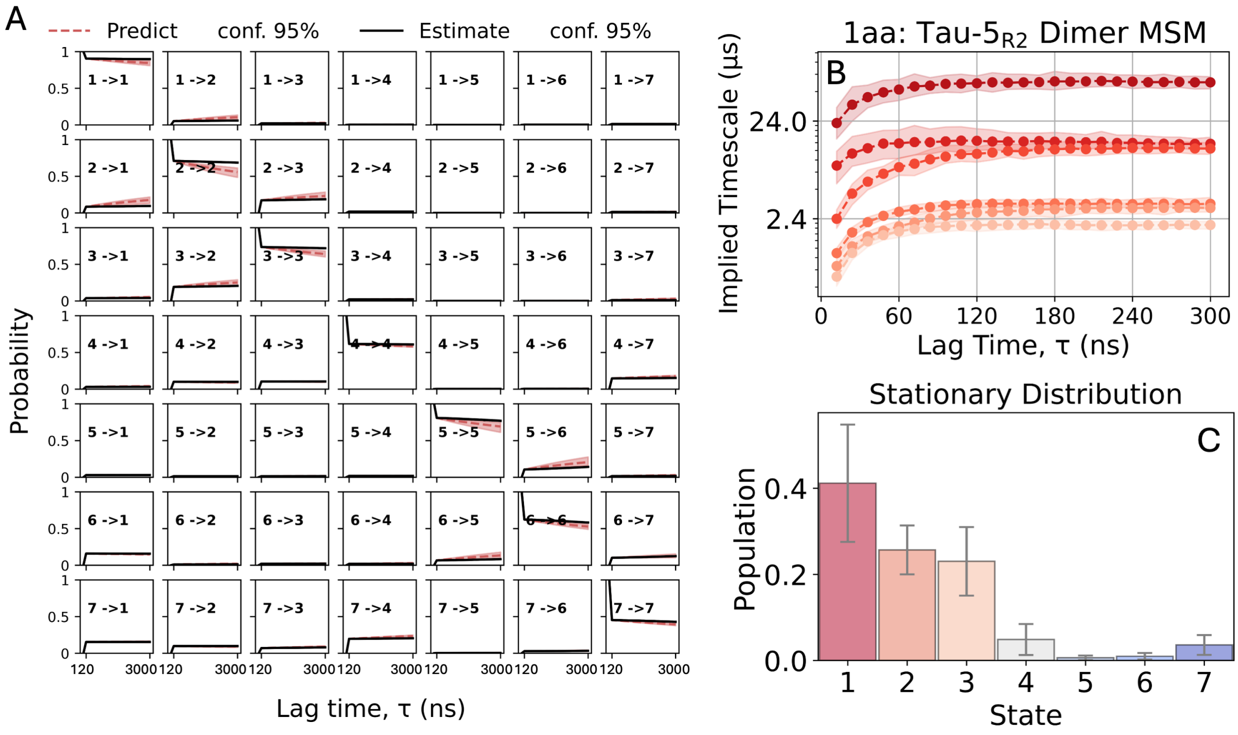


**Supporting Figure S28. Validation tests and stationary distribution for the MSM of Tau-5$_{R2}$ self-association in the presence of 1aa.** (A) Chapman–Kolmogorov (CK) test comparing predicted and directly estimated transition probabilities between states $i$ and $j$ at integer multiples of the model lag time (120 ns). Red dotted lines ("Predict") show transition probabilities predicted by propagating the transition matrix while solid black lines ("Estimate") show transition probabilities obtained from transition matrices estimated directly from the trajectory data at integer multiples of the lag time. Red and black shaded regions indicate the 95% confidence interval of the mean obtained from Gibbs sampling with 1000 samples. (B) Log-scaled implied timescales (ITS) obtained from MSM transition matrices estimated at increasing lag times. Converged implied timescales are expected to be approximately independent of lag time. Colored shaded regions indicate the 95% confidence intervals of the mean implied timescales at each lag time. Here, all ITS are well above the threshold that can be resolved by the model. (C) Stationary probabilities of the MSM states, with error bars indicating the 95% confidence interval of the mean.

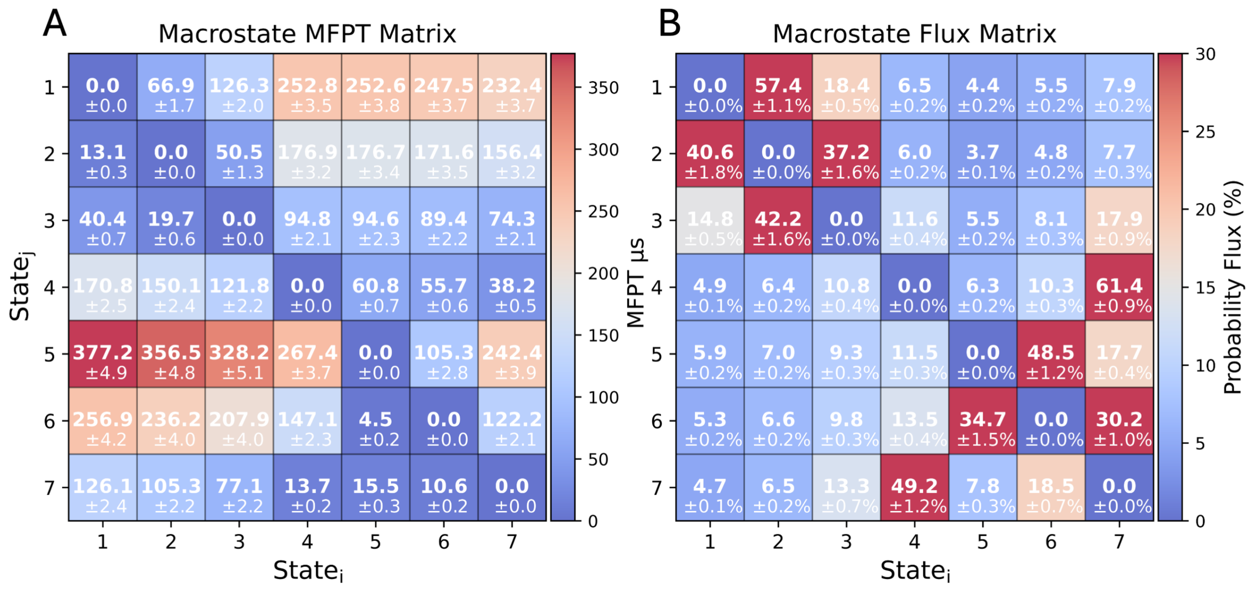


**Supporting Figure S29. Mean first-passage times and transition probability fluxes for the MSM of Tau-5$_{R2}$ self-association in the presence of 1aa.** (A) Mean first-passage times (MFPTs) and (B) row-normalized transition probability fluxes between metastable macrostates obtained by generalized PCCA spectral clustering of an MSM containing 250 microstates.

(15) Harris, C. R.; Millman, K. J.; Van Der Walt, S. J.; Gommers, R.; Virtanen, P.; Cour-

napeau, D.; Wieser, E.; Taylor, J.; Berg, S.; Smith, N. J. et al. Array programming with NumPy. *Nature* **2020**, *585*, 357–362.

(37) Bielskutė, S.; Garcia-Cabau, C.; Frigolé-Vivas, M.; Szulc, E.; De Mol, E.; Pesarrodona, M.; García, J.; Salvatella, X. Low amounts of heavy water increase the phase

separation propensity of a fragment of the androgen receptor activation domain. *Protein Science* **2021**, *30*, 1427–1437.

(38) Shen, Y.; Bax, A. SPARTA+: A Modest Improvement in Empirical NMR Chemical Shift Prediction by Means of an Artificial Neural Network. *Journal of Biomolecular NMR* **2010**, *48*, 13–22.

(39) Cavalli, A.; Montalvao, R. W.; Vendruscolo, M. Using Chemical Shifts to Determine Structural Changes in Proteins upon Complex Formation. *The Journal of Physical Chemistry B* **2011**, *115*, 9491–9494.

(40) Camilloni, C.; De Simone, A.; Vranken, W. F.; Vendruscolo, M. Determination of Secondary Structure Populations in Disordered States of Proteins Using Nuclear Magnetic Resonance Chemical Shifts. *Biochemistry* **2012**, *51*, 2224–2231.
